# Dietary ^14^C reservoir effects and the chronology of prehistoric burials at Sakhtysh, central European Russia

**DOI:** 10.1101/2023.09.04.555011

**Authors:** John Meadows, Anastasia Khramtsova, Elena Kostyleva, Sergey Vasilyev, Elizaveta Veselovskaya, Maria Dobrovolskaya, Nicolas da Silva, Ben Krause-Kyora, Henny Piezonka

## Abstract

We present the first robust radiocarbon (^14^C) chronology for prehistoric burial activity at Sakhtysh, in European Russia, where nearly 180 inhumations attributed to Lyalovo and Volosovo pottery‐using hunter‐gatherer‐fishers represent the largest known mortuary populations of these groups. Past attempts at ^14^C dating were restricted by poor preservation and limited understanding of diet and dietary ^14^C reservoir effects (DREs). We obtained 32 new AMS (Accelerator Mass Spectrometry) ^14^C dates on human petrous bones. Dietary stable isotopes (δ^13^C, δ^15^N) for all AMS‐dated human samples allow us to propose a novel DRE correction model, using differences in ^14^C, δ^13^C and δ^15^N from bones and teeth of the same individuals to predict DREs of up to c.900 ^14^C years. Our chronological model for 40 individuals dates Lyalovo burials to the early 5^th^ millennium cal BC, and Volosovo burials to the mid‐4^th^ to early 3^rd^ millennium. It reveals a previously unrecognised shift in the Volosovo subsistence economy at c.3300 cal BC, coinciding with a reorientation of trade networks, and shows that the last burial at Sakhtysh was the only one in a crouched position, which coincided with the beginning of Fatyanovo practices, the regional expression of the Yamnaya/Corded Ware expansion.

## Introduction

Sakhtysh I, II, IIa and VIII (56° 47’ 08’’ N, 40° 26’ 56” E, Figure 1) are among the best‐known burial grounds of Neolithic (pottery‐using hunter‐gatherer‐fisher, HGF) groups in north‐eastern Europe, comparable in scale to the Neolithic period at Zvejnieki, Latvia [1] and the Mesolithic (aceramic HGF) cemetery at Yuzhniy Oleniy Ostrov, Karelia, Russia [2]. At least 178 individuals were excavated at Sakhtysh between 1962 and 1993 [3,4]. All burials were inhumations, generally of single adults, dated by archaeological evidence to the Lyalovo or Volosovo period (c.5^th^ or c.4^th^ millennium cal BC respectively). Graves were well separated in both periods, with little intercutting. At Sakhtysh IIa, the largest cemetery, c.15 Lyalovo burials were oriented NW‐SE, while c.56 Volosovo burials were mostly on a SW‐NE axis.

**Figure 1.**
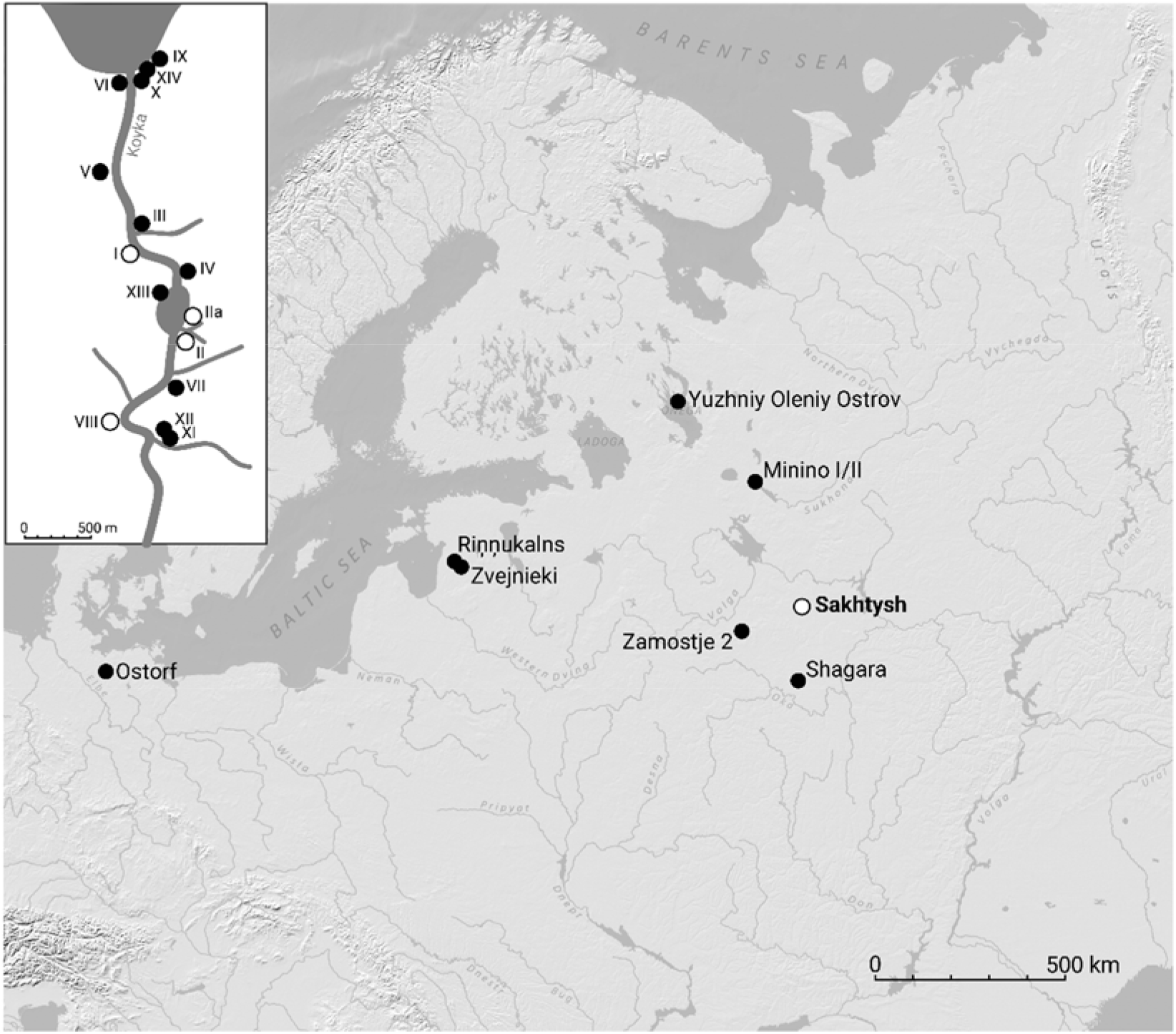
Map of northeastern Europe indicating sites named in the text; inset: the Koyka river valley at Sakhtysh with the locations of prehistoric sites (sampled burial sites highlighted).

Past attempts to date prehistoric burial activity at Sakhtysh have been hampered by poor collagen preservation, inadequate sampling, and inability to quantify dietary ^14^C reservoir effects (DREs) in osseous (bone/tooth) samples. Radiometric ^14^C dating of human bones at Sakhtysh IIa was unsuccessful: 9 samples failed, 8 of 19 reported ^14^C ages were inconsistent with archaeological dating, and even plausible ^14^C ages had uncertainties of at least ±110 years [3]. Piezonka et al. [5] reported the first AMS (Accelerator Mass Spectrometry) ^14^C ages and dietary stable isotopes (δ^13^C, δ^15^N), on bones of 4 individuals from Sakhtysh IIa, and suggested that their high δ^15^N values reflected regular consumption of aquatic animals (fish, invertebrates, water birds; hereafter ‘fish’), leading to DREs (misleadingly old ^14^C ages [e.g., 6]). Bone collagen δ^13^C and δ^15^N of 23 individuals from Sakhtysh IIa [7] supported Piezonka et al. [5]’s suggestion that Lyalovo δ^15^N values were higher than those of Volosovo individuals; in both periods, the range of δ^13^C values was much wider than expected if diets were dominated by terrestrial food sources.

Macāne et al. [8] published AMS ^14^C ages, δ^13^C and δ^15^N values, for 8 animal bones or teeth from Volosovo burials and other ritual contexts at Sakhtysh II and IIa. These should have dated 5 graves, but in one case samples were from an aquatic species, and in another, elk and bear teeth gave inconsistent ^14^C ages. An elk tooth pendant dated previously, which gave an implausibly early date, is regarded as residual [9]. Most burials had no organic grave goods, so accurate dating of human remains is essential, requiring realistic DRE corrections. Macāne et al. [8] noted that freshwater reservoir effects (FREs) may explain the 449±49 ^14^C year difference between 2 food crust samples from the same Volosovo pot, implying that FREs at Sakhtysh were much greater in the past than the c.270 years in 3 fish caught in the river Koyka in 2015 [10]. A preprint of an ancient DNA study [11] includes ^14^C ages on teeth from 18 Sakhtysh individuals, and δ^13^C and δ^15^N for 17 of them. As a guide, Allentoft et al. [11] assume a local FRE of 500 ^14^C years, and that either 50% or 100% of dietary protein came from fish (depending on δ^15^N values), i.e., that individual DREs were 250 or 500 years.

We obtained AMS ^14^C dates and δ^13^C and δ^15^N values on petrous bones of 32 individuals, sampled in 2018. In total, AMS dates are now available for 40 burials in 37 graves. We combine new and existing data in a Bayesian chronological modelling framework to produce a reliable chronology, based on individual DRE estimates. At Sakhtysh, 700 km inland, DREs would have been driven by freshwater fish consumption. We introduce a new approach to freshwater DRE correction, multiple linear regression of ^14^C differences between different skeletal elements of the same individual against δ^13^C and δ^15^N differences in the same samples (‘MLR‐of‐differences’). This approach allows us to address several chronological questions:

- Dating the start, end, and duration of burial activity in each period (Lyalovo, Volosovo)
- Whether Volosovo burials can be split into earlier and later phases
- Whether the Volosovo cemeteries (Sakhtysh I, II, IIa, VIII) were used concurrently
- Whether there was a coherent spatial‐temporal pattern of Volosovo burials at Sakhtysh IIa
- Whether mortuary practices changed over time
- Whether diachronic or synchronic dietary differences between individuals can be detected.

## Results

### New analytical results and legacy data

All but one of our samples produced enough collagen for analysis (Table 1). Carbon and nitrogen contents are close to canonical values for well‐preserved collagen (Supplementary 1) [12‐14], so we assume that AMS and IRMS results are valid.

**Table 1:** new analytical data from Sakhtysh. Each sample consisted of a single human petrous bone. Sex determination M male, F female, U undetermined; upper‐case for genetic sex determination, lower‐case for osteologically determined sex. Chronological attributions based on our dating model.

| laboratory code | cemetery, grave | chronological attribution | sex | age-at-death | % yield | %C | %N | C:N atomic | $\delta^{13}\text{C}$ (‰) | $\delta^{15}\text{N}$ (‰) | <sup>14</sup>C age (BP) |
| --- | --- | --- | --- | --- | --- | --- | --- | --- | --- | --- | --- |
| KIA-56421 | I grave 8 | later Volosovo | M | 40-45 | 7.3 | 40.75 | 14.82 | 3.18 | -20.99 | 13.22 | 4821±28 |
| KIA-53559 | II grave 12 individual A | transitional | M | 30-35 | 4.4 | 41.17 | 14.85 | 3.24 | -22.24 | 12.75 | 4644±24 |
| KIA-53560 | individual B |  | F | 25-35 | 2.9 | 40.36 | 14.73 | 3.20 | -22.32 | 12.87 | 4736±25 |
| KIA-53561 | II grave 15 individual 7 | later Volosovo | F | 25-30 | 7.6 | 41.98 | 15.36 | 3.19 | -23.51 | 13.34 | 5114±26 |
| KIA-53564 | II grave 19 | Lyalovo | U | ? | 9.5 | 41.25 | 15.30 | 3.15 | -20.37 | 12.18 | 6120±29 |
| KIA-53534 | Ila grave 5 | earlier Volosovo | F | 40-45 | 10.9 | 41.20 | 14.88 | 3.23 | -22.55 | 11.56 | 5032±25 |
| KIA-53531 | Ila grave 7A | earlier Volosovo | U | 20-25 | 5.7 | 41.35 | 15.07 | 3.20 | -21.13 | 11.95 | 5147±25 |
| GrM-17396 | Ila grave 7B | earlier Volosovo | F | ~40 | 2.3 | 45.96 | 16.53 | 3.24 | -20.99 | 13.23 | 5145±25 |
| KIA-56422 | Ila grave 9 | later Volosovo | M | 50-55 | 4.0 | 37.87 | 14.02 | 3.21 | -21.93 | 12.64 | 4978±28 |
| GrM-17643 | Ila grave 10 | later Volosovo | F | 20-25 | 2.6 | 41.20 | 14.70 | 3.27 | -23.69 | 12.52 | 4925±30 |
| GrM-17399 | Ila grave 11 | transitional | F | 20-25 | 2.4 | 51.07 | 17.87 | 3.33 | -22.50 | 13.12 | 4730±25 |
| KIA-56423 | Ila grave 13 sk.2 | later Volosovo | F | 50-60 | 5.6 | 40.85 | 15.11 | 3.15 | -23.27 | 12.68 | 4935±28 |
| GrM-17395 | Ila grave 14 | Volosovo | M | c. 40 | 3.7 | 45.11 | 16.25 | 3.24 | -22.43 | 12.09 | 5000±25 |
| KIA-53562 | Ila grave 15 | later Volosovo | M | 30-39? | 6.8 | 42.19 | 15.61 | 3.15 | -24.74 | 14.47 | 5293±26 |
| KIA-54657 | Ila grave 19 | Earlier Volosovo | F | ? | 9.1 | 41.37 | 15.40 | 3.13 | -20.86 | 11.10 | 4931±29 |
| KIA-56424 | Ila grave 22 | (Lyalovo) | f | 20-25 | 0.2 | - | - | - | - | - | - |
| KIA-53532 | Ila grave 25 | earlier Volosovo | F | 30-35 | 5.4 | 40.78 | 14.97 | 3.18 | -23.18 | 11.67 | 5177±25 |
| GrM-17397 | Ila grave 27 | earlier Volosovo | F | 30-35 | 0.5 | 44.98 | 16.30 | 3.22 | -22.23 | 12.03 | 5165±25 |
| GrM-17638 | Ila grave 32 | later Volosovo | M | ? | 1.8 | 43.70 | 15.70 | 3.25 | -22.82 | 12.30 | 4700±25 |
| GrM-17413 | Ila grave 33 | later Volosovo | M | 50-55 | 2.6 | 52.56 | 18.85 | 3.25 | -23.02 | 12.75 | 5030±25 |
| KIA-56425 | Ila grave 35 | Volosovo | m | 35-40 | 5.2 | 41.72 | 15.09 | 3.15 | -23.35 | 13.09 | 5090±28 |
| KIA-53538 | Ila grave 39 | later Volosovo | M | 25-35 | 3.9 | 41.10 | 15.06 | 3.18 | -24.32 | 13.36 | 5218±26 |
| KIA-53539 | Ila grave 40 | Lyalovo | M | 50-60 | 15.1 | 43.25 | 16.05 | 3.14 | -20.84 | 14.42 | 6432±28 |
| KIA-53536 | Ila grave 42 | Lyalovo | M | 20-25 | 12.7 | 39.65 | 14.84 | 3.12 | -21.34 | 14.00 | 6012±26 |
| GrM-17641 | Ila grave 43 | Lyalovo | M | 5-6 | 0.4 | 44.80 | 16.30 | 3.21 | -23.86 | 16.24 | 6755±35 |
| KIA-56426 | Ila grave 46 | later Volosovo | M | 20-25 | 5.1 | 44.77 | 15.96 | 3.23 | -23.27 | 13.72 | 4874±28 |
| KIA-56427 | Ila grave 56 | later Volosovo | M | 25-30 | 4.3 | 42.90 | 15.46 | 3.27 | -23.90 | 13.16 | 4980±28 |
| KIA-53535 | Ila grave 58 | Volosovo | M | 40-45 | 5.1 | 40.57 | 14.70 | 3.22 | -23.47 | 13.90 | 5258±26 |
| GrM-17642 | Ila grave 61 | Lyalovo | F | 20-25 | 2.7 | 43.50 | 15.90 | 3.20 | -22.43 | 15.76 | 6730±30 |
| GrM-17398 | Ila grave 62 | later Volosovo | M | 40-45 | 0.7 | 45.34 | 16.45 | 3.22 | -24.17 | 12.62 | 5035±25 |
| KIA-56428 | Ila grave 66 | later Volosovo | f | 20-25 | 5.2 | 41.86 | 15.43 | 3.16 | -22.61 | 12.86 | 4949±27 |
| KIA-53533 | Ila grave 67 | earlier Volosovo | M | 17-19 | 4.4 | 39.66 | 14.55 | 3.18 | -22.04 | 11.58 | 4967±29 |
| KIA-53563 | VIII 1965 trench 1 grave ? | later Volosovo | F | 30-35 | 7.7 | 41.61 | 15.30 | 3.17 | -23.01 | 13.15 | 5108±26 |

Many of our samples are from individuals also sampled by Kostyleva and Utkin [3] (radiometric ^14^C; n=19); Piezonka et al. [5] (AMS ^14^C, IRMS δ^13^C, δ^15^N; n=4); Engovatova et al. [7] (IRMS δ^13^C, δ^15^N; n=22); and/or Allentoft et al. [11] (AMS ^14^C, n=18, of which IRMS δ^13^C, δ^15^N, n=17) (Supplementary 2). IRMS results from petrous bones and perhaps teeth support earlier claims [5,7] that δ^15^N values are lower in Volosovo than in Lyalovo individuals (Supplementary 3; Figure S2). Mean δ^15^N in Volosovo petrous bones (12.7±0.8‰, n=24) is significantly lower than in Lyalovo petrous bones (14.5±1.6‰, n=5) (heteroscedastic T test, p=0.032). Mean δ^13^C in Volosovo petrous bones is also lower than in Lyalovo samples (‐22.8±1.1‰, against ‐21.8±1.4‰) (homoscedastic T test, p=0.036). Within each period, δ^13^C and δ^15^N are negatively correlated (Lyalovo (10 AMS‐dated samples): r^2^ = 0.702, p_uncorr_ = 0.0024; Volosovo (38 AMS‐dated samples): r^2^ = 0.317; p_uncorr_ 0.0002). These correlations are inevitable if isotopic variation is due mainly to differential consumption of higher‐δ^15^N, lower δ^13^C fish [e.g., 15]. At Sakhtysh, δ^15^N is a good predictor of δ^13^C within each period, but Lyalovo δ^13^C values are 2‐3 ‰ higher than those of Volosovo samples with similar δ^15^N (Figure S3).

Our data confirm that radiometric bone ^14^C ages were often unreliable (Supplementary 4), as assumed by [3] and [8], so we disregard all radiometric dates. AMS ^14^C ages on different skeletal elements of the same individuals are often significantly different [16], by up to 300‐400 ^14^C years (Supplementary 5). There is no pattern of inter‐laboratory offsets, so we assume that results from each AMS laboratory are equally valid. ^14^C differences probably reflect differences in DREs, due to lifetime changes in diet and differences in collagen formation time. Intra‐skeletal ^14^C age differences should thus correspond to δ^13^C and δ^15^N differences, if aquatic and terrestrial foods had different δ^13^C and δ^15^N values. Even if differences <1‰ may be insignificant [e.g., 17], in at least 5 cases diet apparently changed between early childhood (petrous bone) and later childhood/adolescence (tooth). In 4 of these cases, tooth δ^13^C is lower and δ^15^N is higher (Figure 2a), suggesting that weaning [e.g., 18] is not responsible.

**Figure 2:**
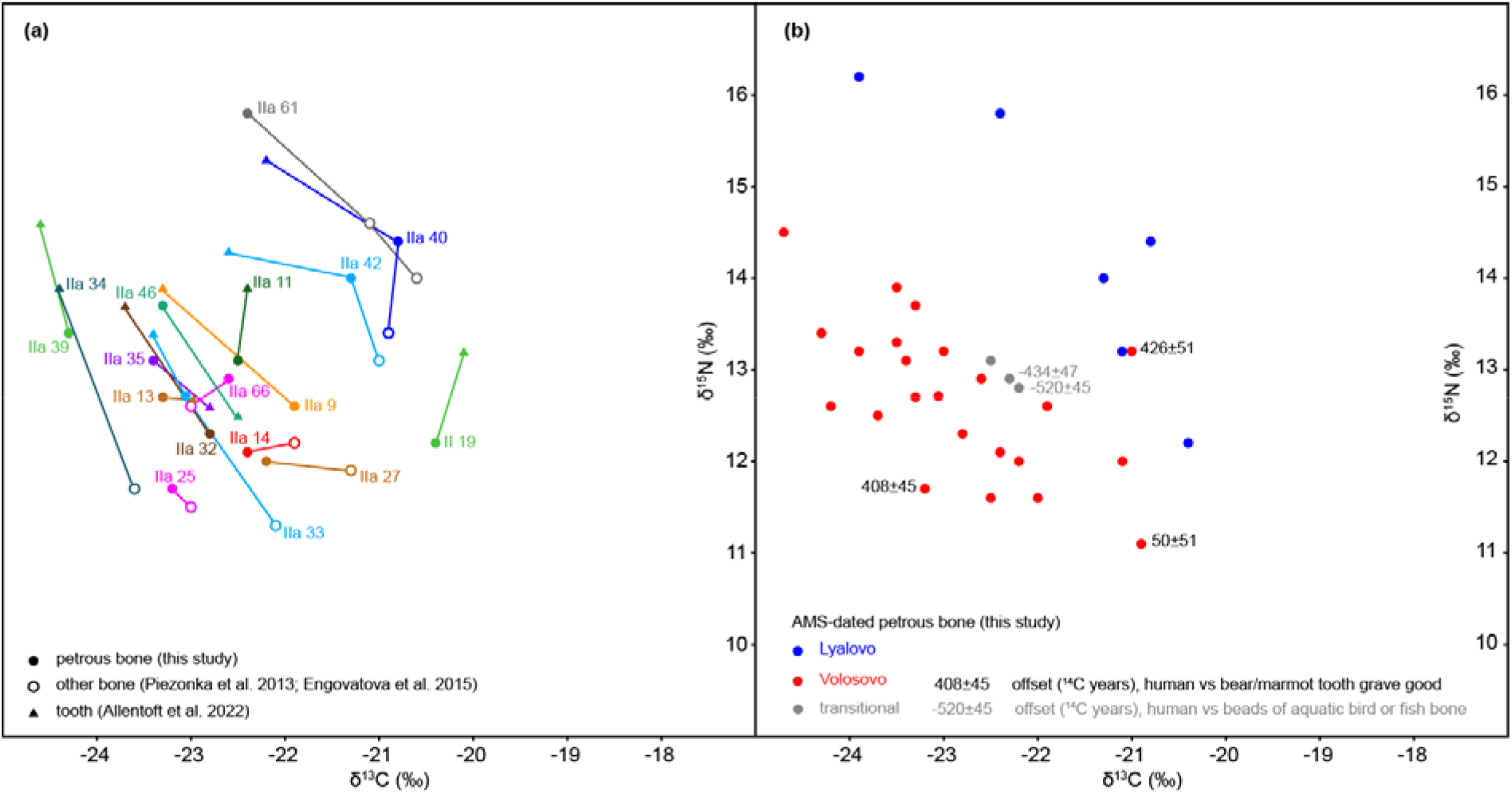
(a) δ^13^C and δ^15^N values from human remains, Sakhtysh prehistoric cemeteries, in cases where we sampled a petrous bone and another skeletal element was analysed by others; lines and colours link results attributed to a single individual (Table 1, Tables S1‐S2); (b) δ^13^C and δ^15^N values from human petrous bones (Table 1), showing ^14^C age offsets in cases where we dated a petrous bone and osseous grave goods were dated by Macāne et al. [8] (Table S4).

We can compare ^14^C ages between human remains and osseous artefacts in only 4 graves (Supplementary 6). The negligible DRE in Sakhtysh IIa grave 19, and c.400 ^14^C year DREs in IIa graves 7 and 24‐25, suggest that δ^13^C and δ^15^N values of e.g. ‐21‰, +11‰ reflect overwhelmingly terrestrial diets, while large DREs are possible if δ^13^C is lower and/or δ^15^N is higher (Figure 2b). Much lower δ^13^C and/or much higher δ^15^N values suggest that in some cases DREs were substantially greater than 400‐500 ^14^C years. The II grave 12 bone beads were from an aquatic species [8], with reservoir effects apparently greater than DREs in the corresponding human samples (Table S3).

### Multiple linear regression of ^14^C age, δ^13^C and δ^15^N differences

There are too few paired human‐herbivore samples to express DREs as a function of human δ^13^C and δ^15^N [19] (see Methods), but in 15 cases we can compare ^14^C age, δ^13^C and δ^15^N between different skeletal elements of single individuals. We can also compare DREs, δ^13^C and δ^15^N in burials IIa 7B and IIa 25 to those of IIa grave 19 (Supplementary 7). In these 17 cases, lower δ^13^C is associated with higher δ^15^N (Figure 3a) (r^2^=0.299, p_uncorr_=0.023). Lower δ^13^C and higher δ^15^N are both apparently associated with higher ^14^C ages (Figure 3b,c), as expected if differential fish consumption causes variation in δ^13^C, δ^15^N, and ^14^C ages.

**Figure 3:**
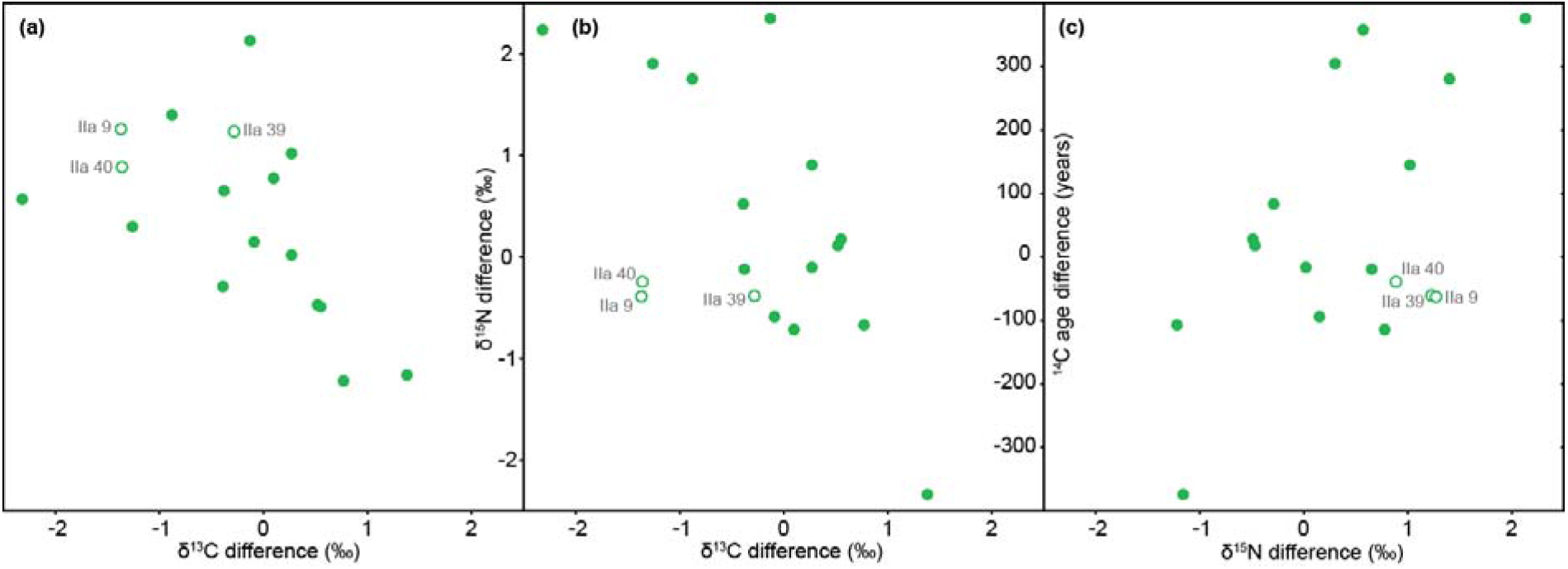
Intra‐skeletal differences in (a) δ^13^C and δ^15^N, (b) δ^13^C and ^14^C age, (c) δ^15^N and ^14^C age, in 17 cases where two samples of the same individual have been AMS‐dated. Circles denote the 3 cases omitted from the regression.

Multiple linear regression (MLR) of all 17 cases shows that δ^13^C and δ^15^N differences explain only about 33% of variation in ^14^C differences, however. IIa graves 9 and 40 (Volosovo and Lyalovo respectively) are outliers (Figure 3b,c). Omitting these 2, MLR accounts for 61% of intra‐skeletal ^14^C variation; ^14^C differences are correlated with δ^13^C differences (r^2^=0.590; p_uncorr_=0.011) but the δ^15^N correlation (r^2^=0.394; p_uncorr_=0.126) could still have arisen by chance (i.e. p_uncorr_>5%). If we also omit IIa grave 39 (Volosovo), 68% of intra‐skeletal ^14^C variation is explained by MLR, and both δ^13^C and δ^15^N are informative. The slope of the 14‐case regression is ‐130±41 for δ^13^C (r^2^=0.623; p_uncorr_=0.009) and 93±40 for δ^15^N (r^2^=0.505; p_uncorr_=0.042), i.e. ^14^C age increases by 130±41 years for every 1‰ decrease in δ^13^C, and by 93±40 years for every 1‰ increase in δ^15^N.

This formula predicts DREs in human samples as a function of differences between their δ^13^C and δ^15^N, and δ^13^C and δ^15^N for a 100% terrestrial diet, which we estimate as ‐20.0‰ and 11.6‰ respectively, based on faunal data (below). The formula accurately predicts DREs in the 3 cases where they are known (Supplementary 8), and predicts much larger DREs in samples with lower δ^13^C and/or higher δ^15^N. These estimates appear realistic, however. For example, IIa grave 32’s tooth (predicted DRE 704±180 years) has a ^14^C age c.300 years greater than that of its petrous bone (predicted DRE 457±126 years), which is isotopically similar to the IIa 25 petrous bone (observed DRE 426±51). The formula predicts DREs of c. 400 years for both individuals in II grave 12, which, given the ^14^C ages of the accompanying aquatic bone beads (Table S4), suggests an FRE in local fish of c.900 years, large enough to cover the range of predicted DREs, provided that in some cases, most dietary protein came from fish.

### Diet reconstruction

We apply quantitative diet reconstruction using FRUITS [20] to check if such diets are consistent with stable isotope results, focussing on dietary protein, which determines δ^13^C and δ^15^N in human collagen, particularly in high‐protein diets. Our FRUITS model (see Methods) only considers terrestrial herbivore and fish protein sources, whose average δ^13^C and δ^15^N can be inferred from archaeozoological data. Plant foods will have been too similar isotopically to terrestrial animals, and too low in protein, to be treated as a separate protein source. The 8 dated animal bones and teeth from Sakhtysh were analysed by EA‐IRMS [5,8], but these finds were portable artefacts, which may be non‐local, and 4 were from bears. Average δ^13^C and δ^15^N for 64 prehistoric herbivores from the region, including the Sakhtysh elk teeth (mean δ^13^C ‐21.5±0.4‰, δ^15^N 5.1±0.2‰, n=2), are ‐22.0±0.5‰ and 5.1±1.2‰ (Supplementary 9). The fish situation is more complex, as fish bones from Sakhtysh IIa did not yield collagen, and regional sites show contrasting patterns (Supplementary 9), which may not account for human data at Sakhtysh. Fish bones from 4^th^‐millennium Riņņukalns, Latvia [21] suggest that an average fish collagen δ^13^C of ‐27±1‰ and δ^15^N of 9.0±1‰ during the Volosovo period may be realistic. Average δ^13^C and/or δ^15^N in fish consumed in the Lyalovo period must have been higher (see Discussion).

Whatever isotope values are applied to Lyalovo fish collagen, FRUITS output supports the impression that diets varied widely between individuals and during individual lifetimes. FRUITS output confirms that some Volosovo diets were based mainly on terrestrial protein (presumably hunted herbivores, such as elk and beaver), while others relied almost entirely on fish (and perhaps aquatic birds and invertebrates). Sakhtysh IIa grave 19 had a mainly terrestrial diet (16.6±8.5% aquatic protein), while IIa grave 15 (88.5±7.2% aquatic protein) represents the aquatic end of the spectrum. However, we cannot distinguish groups of hunters and fishers: mixed diets, in which fish provided 50‐70% of dietary protein, appear to predominate numerically. Mixed diets might represent longer‐term averages than the more specialised extremes, as collagen formation time varies between samples and is unknown in individual cases, but in this case we should not find diachronic patterns in Volosovo diets.

### Bayesian chronological modelling

Our OxCal model (see Methods, Supplementary 10) has good overall agreement (A_overall_ = 90; A_model_ = 121; a threshold of 60 is acceptable [22]), because DRE‐corrected dates of multiple samples from the same burial are mostly compatible. This is not a rigorous test, due to large uncertainties in predicted DREs, but the MLR‐of‐differences formula does not over‐estimate DREs in burials with dated osseous grave goods. Predicted DREs appear to be under‐estimated in IIa grave 7, perhaps because the ^14^C age of the associated bear tooth is slightly too recent. The OxCal output favours an 800‐year FRE for the bone beads in II grave 12. This grave, and the crouched burial IIa grave 11, are not regarded as Volosovo burials in the model, which dates them to the end of the Volosovo period, or slightly later, another indication that the chronology obtained is realistic.

## Discussion

### The long chronology of burial at Sakhtysh

Only 5 Lyalovo burials have been dated, with large uncertainties in their DRE‐corrected dates. Our model dates Lyalovo burial activity to the earlier 5^th^ millennium cal BC (Figure 4a). No burials are recorded between the mid‐ 5^th^ millennium and the first Volosovo burial, in the 37^th^ century cal BC. There are no known later Lyalovo or earlier Volosovo cemeteries in the region corresponding to this hiatus, but ^14^C dating of isolated burials such as Berendeyevo 1 [23](see below) may eventually fill the gap [4]. At Sakhtysh, a single ^14^C age (AAR‐21042; [9]) dates an artefact redeposited in a Volosovo grave to the late 5^th^ millennium, and an elk tooth from Sakhtysh II burial 18 [8] could date to the earlier 4^th^ millennium, just before the first Volosovo burials and ritual deposits. The latest burials took place in the early 3^rd^ millennium, with the last regular Volosovo burial in c. 2900 cal BC. This may have coincided with Sakhtysh II grave 12, which we have treated as culturally transitional on account of its anomalous mortuary style. The latest‐dated burial, IIa grave 11, was originally attributed to Lyalovo [3], but unusually was in a crouched position, which is typical of (Corded Ware‐related) Fatyanovo burials of the mid‐3^rd^ millennium [11]. It apparently postdates both the last Volosovo burial and II grave 12, and could coincide with the start of Fatyanovo burial at other sites in the region [23,24].

**Figure 4.**
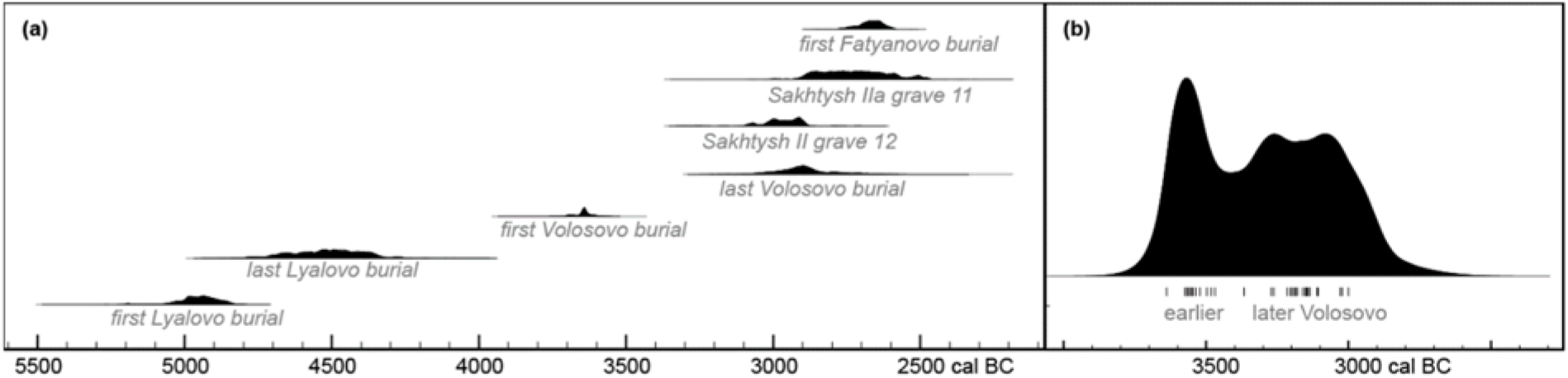
(a) modelled dates of the first‐ and last‐dated Lyalovo and Volosovo burials at Sakhtysh (OxCal functions First and Last) and of transitional burials, II grave 12 and IIa grave 11; our model of 26 human bone calibrated dates from 12 sites in central Russia [data from 23]) (Supplementary Information) dates the first Fatyanovo burial to the 27th century cal BC; (b) kernel density estimate (OxCal function KDE_Model [56]) of the temporal distribution of DRE‐corrected calibrated dates of Volosovo burials at Sakhtysh. Lines show the median modelled dates of individual burials.

Assuming that the temporal distribution of dated burials is representative, there were two peaks of Volosovo burial activity, at c.3500 and c.3000 cal BC (Figure 4b). We refer to 13 mid‐4^th^ millennium burials as earlier Volosovo, and to 16 late 4^th^‐early 3^rd^ millennium burials as later Volosovo. Few burials (IIa graves 14, 35 and 58) are not clearly attributed to one phase (Supplementary 11). A brief hiatus at c.3400‐3300 cal BC is possible.

Most of the 32 AMS‐dated Volosovo burials are from Sakhtysh IIa. The others are Sakhtysh I grave 8 and an individual from Sakhtysh VIII, both dating to the last third of the 4^th^ millennium, and 3 burials at Sakhtysh II. A human tooth from Sakhtysh II grave 4 and a bear tooth from grave 18 date them to the mid‐4^th^ millennium, like Sakhtysh II hoard 9, dated by a bear tooth [8]. Sakhtysh II grave 15 (a multiple burial) dates to the late 4^th^ millennium, like the badger bone from hoard 11 [8]. The anomalous II grave 12 multiple burial is one of the latest graves at Sakhtysh, dating to the early 3^rd^ millennium. Thus the other cemeteries do not fill potential gaps in burial activity at Sakhtysh IIa, either during the early 4^th^ millennium or at c.3400‐3300 cal BC.

Spatial‐temporal patterning helps to explain the lack of intercutting graves at Sakhtysh IIa, despite the extended period of burial. Several rows (NW‐SE alignments of parallel graves) were recognised. Based on grave depths, [3] proposed that rows A, B (Б) and Е predated rows V (B), G (Г), D (Д) and Zh (Ж), but all 3 dated burials in Row G (Г) are earlier Volosovo, while all 7 dated burials in rows A and E are from the later Volosovo phase. In general, earlier Volosovo burials are concentrated in rows G (Г) and B (Б), but the later Volosovo burial in grave 36 (at the SE end of row D (Д)) cut an earlier Volosovo burial. Graves 63 and 67 at the NW edge of the cemetery are also earlier Volosovo (Figure 5), as is grave 54 in the ‘sanctuary’ area. Later Volosovo burials are concentrated on the north edge of the sanctuary (rows D (Д), E) and the south edge of the cemetery (row A).

**Figure 5.**
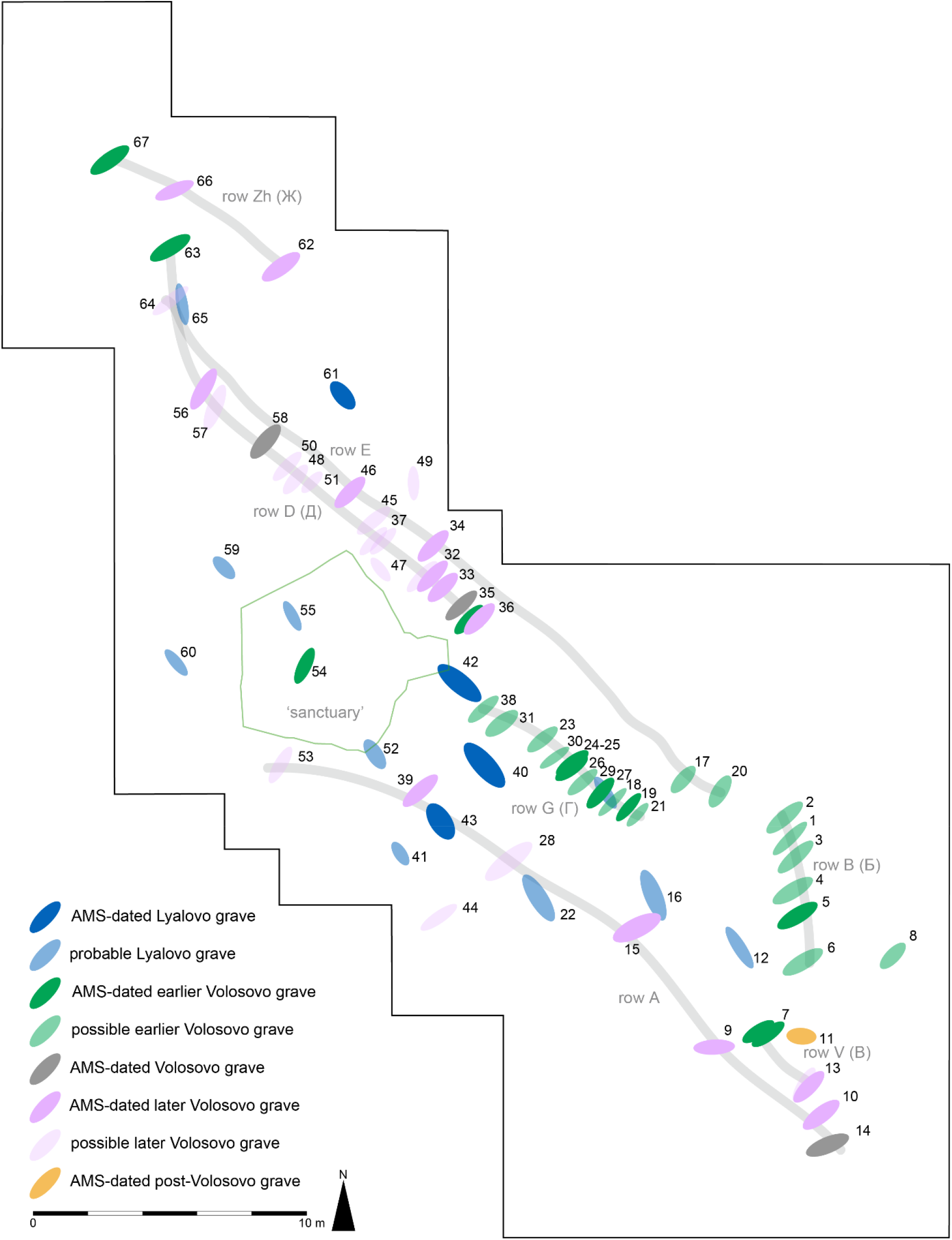
Attribution of Sakhtysh IIa graves to chronological phases. Bold colours denote AMS‐dated burials, phasing based on our DRE corrections and chronological model. Pale colours indicate the most likely phasing of graves without AMS dates, based on grave orientation and location.

Sampling was not targeted at grave goods, but some chronological patterns emerge (Figure 6a). Bear teeth are practically restricted to earlier Volosovo burials, which is why Macāne et al. [8] proposed a much shorter period of Volosovo burial (‘tentatively 3650–3400 cal BCE’). Osseous grave goods in general are far more common in earlier graves, and of 8 dated burials with slate and/or serpentine, derived from the Urals, 6 date to the earlier Volosovo phase. All 4 dated cases with Baltic amber date to the later Volosovo‐transitional phases, suggesting a shift in the orientation of trade networks in c.3400‐3300 cal BC.

**Figure 6:**
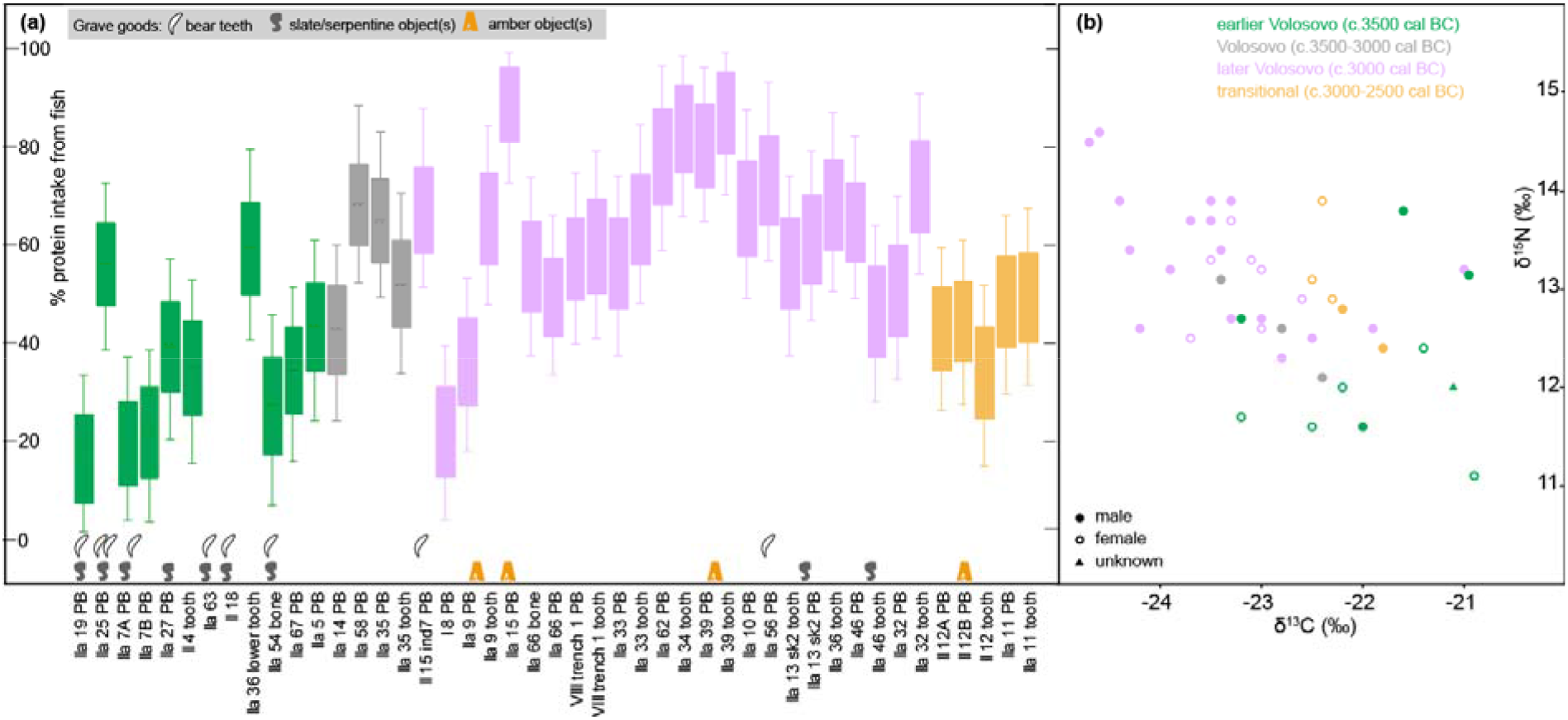
(a) grave goods accompanying dated burials, in median date order (Bayesian chronological model output), and FRUITS box‐and‐whisker (68%/95% probability) estimates of fish protein intake in AMS‐dated Volosovo‐transitional human samples from Sakhtysh; (b) δ^13^C and δ^15^N in these samples (symbols indicate sex and phasing).

### Wide variation in Volosovo diets is partly a diachronic pattern

Lyalovo δ^13^C and/or δ^15^N values are consistently higher than those of Volosovo samples, but the range of values is similar in each phase (Figure S3). If isotopic signatures of potential foods were the same in each period, diet‐ collagen fractionation (for δ^13^C, δ^15^N or both) might have been greater in Lyalovo individuals, due to unknown metabolic differences (any difference in weaning age is irrelevant, because we see the same offset in adult bones as in teeth and petrous bones). More plausibly, Lyalovo fishers may have targeted only higher trophic‐level fish, while Volosovo fishers used a wider range of species. Alternatively, environmental change may have increased access to e.g. freshwater molluscs, and/or reduced access to fish with higher δ^13^C. A combination of factors may apply, but a shift in the average δ^13^C (and perhaps δ^15^N) of fish consumed seems the most realistic explanation for the Lyalovo‐Volosovo isotopic offset (Supplementary 12).

We use FRUITS [20] to convert human δ^13^C and δ^15^N to diets, which requires several assumptions (see Methods). Most pertinently, we assume that collagen δ^13^C and δ^15^N are only affected by dietary protein, which means that FRUITS only quantifies protein sources and effectively disregards plant foods. In reality, it is unlikely that boreal HGFs would have survived without energy‐dense foods, which are barely visible in collagen δ^13^C and δ^15^N. For simplicity, we assume the same average δ^13^C and δ^15^N in fish consumed by each individual within each period, even if average fish δ^13^C (and/or δ^15^N) was apparently higher in the Lyalovo period. Significant correlations between human δ^13^C and δ^15^N within each period support this assumption.

Not enough Lyalovo individuals have been dated to reveal diachronic or synchronic dietary patterns, beyond the observation that diets seem to have varied widely between individuals (fish providing c.20% to c.95% of protein intake) and could change dramatically during one lifetime. Among Volosovo individuals, any age‐related dietary patterns are hidden, as most AMS‐dated samples were formed in childhood‐adolescence, and adult bone δ^13^C and δ^15^N [7] may not be comparable (Supplementary 3). δ^13^C or δ^15^N differences between AMS‐dated samples from Volosovo males (n=23) and females (n=14) are not statistically significant. Later Volosovo males had the most aquatic diets, but males are over‐represented in the later Volosovo and females are over‐represented among earlier Volosovo samples. In both sexes, δ^13^C is visibly lower and δ^15^N higher in later Volosovo than in earlier Volosovo samples; samples that could date to either phase have intermediate values (Figure 6b).

Mid‐4^th^ millennium individuals had mainly terrestrial diets (fish providing about a third of dietary protein), a pattern reflected in the earlier Volosovo dates of osseous grave goods [8] (Figure 6a). Most late‐4^th^ millennium individuals had much more aquatic diets (typically two‐thirds fish protein), comparable to those of contemporaneous HGFs at Lake Burtnieks, Latvia [15,25], and Ostorf, Germany [26]. Extreme reliance on fishing might thus be a widespread late 4^th^ millennium adaptation. However, increased fish consumption over time at Sakhtysh also fits the general trend in long‐lived HGF cemeteries in the Lake Baikal region of Siberia, such as the contemporaneous Isakovo cemetery at Ust’‐Ida I [27], which has been attributed to over‐hunting. Terrestrial herbivores perhaps learned to avoid areas with concentrations of HGFs, leading to increased reliance on fishing, which reduced HGF mobility, reinforcing the trend. If this is a realistic model for Siberia, it might also apply at Sakhtysh.

The latest burials (Sakhtysh II grave 12 (both individuals) and IIa grave 11) had more mixed diets, consuming similar amounts of terrestrial and aquatic protein (Figure 6a). Burial rites in both graves are anomalous for the Volosovo period and IIa grave 11’s crouched position suggests contact with Fatyanovo culture [11], in which pastoralism may already have been established [28].

Synchronic dietary variation is harder to assess, due to uncertainties in DRE‐corrected burial dates, but lifetime diet changes are indicated by δ^13^C and δ^15^N differences between petrous bones and teeth of single individuals. These shifts (up to 1.4 ‰ in both isotopes) are much smaller than the overall δ^13^C and δ^15^N ranges in each period (c.4 ‰ in both isotopes, c.3 ‰ in just the later Volosovo phase), suggesting that there were persistent differences in personal dietary preferences among broadly contemporaneous individuals.

In a regional context, there are no close palaeodietary analogies for the Lyalovo cases. A late 5^th^ millennium date for a single individual from Berendeyevo, only 90 km west of Sakhtysh, was published without δ^13^C and δ^15^N [23]. The two individuals dated to c.5000 cal BC at Minino I, 330 km north of Sakhtysh, have δ^13^C and δ^15^N values consistent with a primarily fish‐based diet, and large DREs [6]. Otherwise we have to consider Zvejnieki, Latvia, 920 km west of Sakhtysh, where in the corresponding period there appears to have been a wide range of individual diets along a gradient from mainly terrestrial to mainly based on local fish [15]. Human δ^13^C and δ^15^N values at Neolithic HGF cemeteries beside the Dnipro Rapids, Ukraine, 1000 km south of Sakhtysh, demonstrate more consistently fish‐based diets than in earlier or later periods, although accurate dating of individual burials remains problematic [29‐32]. By the 4^th^ millennium BC, however, this region had a farming or pastoral economy. The most comparable data for Volosovo burials are from Shagara, only 175 km south of Sakhtysh, where 19 HGF individuals attributed archaeologically to the Volosovo period were analysed [33]. Quantitative diet reconstruction [33] suggested that fish provided only 25±14 % of protein in Shagara Volosovo diets, much less than in the later Volosovo population at Sakhtysh, but comparable to earlier Volosovo burials; ^14^C dates of the Shagara burials are unpublished.

### A new approach to freshwater DRE correction in human bones

Ideally, freshwater DREs can be estimated by scaling ^14^C offsets in paired samples of human remains and contemporaneous terrestrial materials, such as bone grave goods, to human dietary stable isotope values [19,34,35]. The diet‐reconstruction approach to freshwater DRE correction has been used when paired terrestrial samples are not available, but applicable FRE and food‐source stable isotope values are understood [e.g., 15,36]. We could not apply either approach at Sakhtysh because bone grave‐goods were not accessible and the FRE in local fish was unknown. We initially used a FRUITS diet‐reconstruction model to estimate fish contributions to δ^13^C and ^14^C age in AMS‐dated human samples, and then Bayesian chronological modelling to find the range of FRE values required to reconcile human and grave‐good ^14^C ages, and ^14^C ages of multiple samples from single individuals. This exercise suggested an average FRE in local fish of at least 900 years. FREs of this magnitude are not unusual in north‐eastern Europe [e.g., 36,37; Meadows unpublished data].

The largest DREs predicted by our 14‐case MLR‐of‐differences model (965±252, 912±233, 906±231 years) are of a similar order, and our FRUITS model suggests that fish provided nearly all dietary protein in these cases. The principle that the MLR‐of‐differences and diet‐reconstruction approaches should give similar DRE estimates if realistic parameter values are applied allows some sensitivity analysis of these values (Supplementary 13). We cannot test whether our expected δ^13^C and δ^15^N values for 100% terrestrial diets are valid, as the same values correspond to zero DRE in either model. Changing them would systematically shift DREs (by 9‐13 years for every 0.1 ‰ in the MLR‐of‐differences model). Using our preferred fish δ^13^C and δ^15^N values in the FRUITS model, a 960‐year average FRE leads to consistent DRE estimates for all Volosovo‐transitional burials, including those not used in the MLR‐of‐differences model, but Lyalovo diet‐reconstruction DRE estimates are too low (Figure S7). Unsurprisingly, fish δ^13^C and δ^15^N values giving valid estimates of fish intake in Volosovo cases appear to under‐ estimate Lyalovo fish consumption. The MLR‐of‐differences approach, which relies on δ^13^C and δ^15^N differences within individuals, should be less sensitive to baseline shifts between periods in δ^13^C and/or δ^15^N than DRE correction based on ^14^C offsets in paired human and terrestrial samples.

Even a MLR approach using ^14^C offsets in paired human‐terrestrial samples will encounter outliers (e.g. non‐local individuals), which may be omitted from the regression. The presence of outliers among perfectly‐paired cases implies that some of the regression‐based DRE estimates for unpaired individuals will be misleading (e.g. if an unpaired individual was non‐local). Whether this matters depends on the chronological questions, and on how many DRE estimates are misleading. After removing outliers, r^2^ values >80% are achieved in some of the Baikal case studies [19]. A lower r^2^ is to be expected in an MLR‐of‐differences regression, as measurement uncertainties are proportionally larger, so the data must be noisier. At Sakhtysh, we omitted 3 of the 17 potential cases, improving adjusted r^2^ from 33% to 68%; at this point, correlations between ^14^C differences and δ^13^C and δ^15^N differences cannot have arisen by chance (p_uncorr_<5% for both isotopes). A higher r^2^ and more precise DRE predictions can be obtained by omitting more cases, but this would imply that more of the predicted DREs are misleading. The 3 cases omitted from our MLR‐of‐differences model are not outliers in the δ^13^C vs δ^15^N difference plot (Figure 3a). This suggests that one of the ^14^C ages in these cases is an outlier, perhaps because these individuals moved around more than the others. If the 17 cases with multiple AMS dates are representative, c.9 of the 53 predicted DREs in our chronological model may be misleading. OxCal output identifies 3 cases in which DRE‐corrected dates from the same burial are slightly inconsistent, but the combined burial date estimates are probably reasonable. The issue is how a few cases with one AMS date and a misleading predicted DRE would affect the site chronology. At Sakhtysh, DRE‐corrected dates span several centuries, and serve to attribute burials to phases, not to provide a high‐resolution chronology, so the impact of occasionally misleading DRE estimates should be limited.

## Conclusion

Earlier attempts to date prehistoric human remains at Sakhtysh were frustrated by limited sampling and unreliable analytical results, but beyond these issues, inability to quantify DREs meant that the absolute chronology of these cemeteries was inaccessible. We have addressed this problem by using intra‐skeletal differences in isotopic data to create a DRE correction formula. This shows that, despite large uncertainties in the dates of individual burials, there was clearly a long hiatus between the last Lyalovo burial, in the mid‐5^th^ millennium cal BC, and first Volosovo burial at Sakhtysh in c.3600 cal BC. Volosovo burials span 6‐7 centuries, until c.2900 cal BC. This period can be divided into an earlier Volosovo phase, apparently characterised by more terrestrial diets and grave goods, with connections to the east, and a later Volosovo phase, with more fish‐based diets and westward connections. Before AMS dating, it was assumed that burials with amber ornaments belonged to the earlier Volosovo phase, and those with serpentine to its later phase. Thus our chronological model requires a fundamental revision of the previous periodization of Volosovo culture, and by reversing the traditional sequence, allows us to look afresh at the problem of the origin of the Volosovo culture. With AMS dating, but without the large DRE corrections predicted by MLR‐of‐differences our model, these two phases would appear to be contemporaneous, and what was probably an important diachronic change would be indistinguishable from synchronic variation. Our model dates the only crouched burial to soon after the end of the Volosovo period, coinciding with the start of crouched burial practice with the appearance of Fatyanovo burials in this region. The Sakhtysh case study demonstrates both the value of accurate DRE correction, and that when independent dating of burials is not accessible, realistic estimates of DREs can be obtained by comparing isotopic signals from different elements of the same skeletons.

## Methods

### Laboratory methods

This study is based on new Accelerator Mass Spectrometry (AMS) and Elemental Analysis‐Isotope Ratio Mass Spectrometry (EA‐IRMS) results on collagen extracted from human petrous bones, obtained from 32 of the 62 individuals available for ancient DNA sampling [38]. Petrous bones were sampled because they are usually well preserved, providing good collagen yields and DNA preservation.

Collagen was extracted by the AMS laboratories in Kiel and Groningen following standard protocols [39,40]. At room temperature, ∼1 g of crushed bone fragments were demineralized in HCl, treated with NaOH to dissolve secondary organic compounds, and re‐acidified in HCl, before gelatinization overnight in a hot (75–85 °C) pH3 solution, and filtration to remove insoluble particles. Collagen extracts were freeze‐dried and weighed to determine yield as a percentage of the starting weight.

A sufficient quantity of each extract was combusted, and the CO_2_ obtained was reduced to graphite for AMS measurement. Kiel used a 3 MV HVEE Tandetron AMS, in operation since 1995 [41] and upgraded in 2015. Groningen used a 180 kV IonPlus Micadas AMS system, installed in 2017. Both systems measure ^12^C, ^13^C and ^14^C currents simultaneously; the ^13^C/^12^C ratio (AMS δ^13^C) is used to normalise the ^14^C current for natural and instrumental fractionation, and thus to obtain conventional ^14^C ages [42]. The reported ^14^C age errors incorporate uncertainties in measurement, standard normalisation, instrumental background, blank correction, and additional uncertainty arising from sample pretreatment, based on long‐term experience with laboratory standard and known‐age samples of similar materials [43].

Stable isotope results are expressed using δ notation (δ = [(R_sample_/R_standard_‐1)]×1000, and R = ^13^C/^12^C or ^15^N/^14^N) in parts per mille (‰) relative to international standards, Vienna PeeDee Belemnite for δ^13^C, and air N_2_ for δ^15^N. At Groningen, collagen was combusted in an EA (measuring %C, %N) before graphitisation, with part of the resulting gas directed to an IRMS for δ^13^C and δ^15^N measurement, with estimated uncertainties of ±0.15‰ and ±0.3‰ respectively [40]. Leftover collagen from samples extracted in Kiel was sent for EA‐IRMS to isolab GmbH, Schweitenkirchen, Germany, for measurement of %C, %N, %S, δ^13^C, δ^15^N and δ^34^S [44]. Four aliquots of each sample were analysed, with final measurement uncertainties better than ±0.1‰ for δ^13^C and δ^15^N. Sulfur results are not discussed here due to a lack of reference data or comparable studies in this region, but do not alter our interpretations. In the 22 cases for which we have δ^34^S data, quality assurance criteria are satisfied, but there is no apparent correlation between δ^34^S and either δ^13^C or δ^15^N. This pattern suggests that terrestrial and aquatic fauna δ^34^S values at Sakhtysh overlapped and that δ^34^S would be uninformative even if it could be used in the MLR‐of‐differences model.

### Modelling tools

Two approaches to freshwater DRE correction are in use: mathematical fitting of ^14^C offsets between human bone and contemporaneous terrestrial organisms to human dietary stable isotope values (‘perfect pairs’) [2,19,34,45], and quantitative diet reconstruction, combined with independent estimates of FREs in aquatic species [26]. Where FREs are well‐constrained, and isotope values of potential food sources are understood, these approaches should provide compatible and credible DRE corrections.

Here, we introduce a new version of the first approach, multiple linear regression of ^14^C offsets between different skeletal elements of the same individual against differences in δ^13^C and δ^15^N values from the same samples (‘MLR‐of‐differences’), incorporating ^14^C offsets relative to terrestrial animal teeth in the 3 cases where this information is available. We use Past 4.10 (https://www.nhm.uio.no/english/research/infrastructure/past/) [46] for simple graphics and statistical analyses of new and published ^14^C, δ^13^C and δ^15^N values, including multiple linear regression.

We also apply the diet reconstruction approach, using the Bayesian statistical package FRUITS [20] and published faunal δ^13^C and δ^15^N values from prehistoric sites [33,47], and modern data [48] from the same region. Expert inspection of scatter plots of δ^13^C and δ^15^N data often provides insights into the importance of potential food sources, but it is hard to quantify the intake of different foods, and uncertainty in such estimates, without a formal statistical model. For our purposes, one advantage of FRUITS over alternatives, e.g. MixSIAR (https://cran.r-project.org/web/packages/MixSIAR/index.html), is that the model output includes posterior estimates of the contribution of each food group to each isotope value. Since the contribution of fish to δ^13^C and ^14^C must be identical, FRUITS posterior estimates of fish contribution to δ^13^C can be used to predict DREs, based on potential FREs in the fish consumed. We test a range of potential FREs to find which FRE values give compatible DRE predictions to those given by the MLR‐of‐differences formula.

Our FRUITS diet reconstructions only address dietary protein sources, which are assumed to determine collagen δ^13^C and δ^15^N values, although energy macronutrients (fats and carbohydrates) can have a measurable impact on collagen δ^13^C values, particularly in low‐protein diets [49]. HGFs in boreal forest environments probably had high‐protein diets, however, and an unrouted, protein‐only model [50] appears to provide more accurate estimates of the effect of fish consumption on collagen stable isotope values and ^14^C ages [51]. This assumption simplifies the modelling, and is less critical than other necessary assumptions. Terrestrial food sources include both plants and animal tissues (but not dairy at this date), but flesh and blood would have contained much more protein than plant foods, so the isotopic signature of terrestrial foods can be inferred from animal collagen δ^13^C and δ^15^N. Based on data from modern animals [e.g., 50], we assume that flesh protein had δ^13^C 2.5‰ lower and δ^15^N 1.5‰ higher than faunal bone collagen, regardless of species. The isotopic spacing between dietary protein and human collagen cannot be measured without long‐term controlled feeding studies, and estimates of ^15^N fractionation vary considerably (c.+3 to +6‰). We apply diet‐collagen offsets of +4.5±0.25‰ (i.e. +4 to +5 ‰) for δ^13^C and +5.0±0.5‰ (i.e. +4 to +6 ‰) for δ^15^N. Reducing the δ^15^N offset would increase the estimated intake of higher‐δ^15^N fish protein. At Sakhtysh, most of the isotope data are from petrous bones or teeth, creating another uncertainty: to what extent these results incorporate a ‘nursing effect’, i.e. whether some of the collagen was formed in early infancy when the child was breastfed, and thus at a higher trophic level. However, higher δ^15^N is usually associated with lower δ^13^C, which can be explained by fish consumption but not by nursing.

For chronological modelling, we use OxCal v4.4 (https://c14.arch.ox.ac.uk/oxcal/OxCal.html) [e.g., 22]. Estimated DREs can be subtracted arithmetically from uncalibrated ^14^C ages before calibration with the relevant atmospheric calibration curve, currently IntCal20 [e.g., 2,26,50,52]. Alternatively, in OxCal, estimated DREs can be applied to uncalibrated ^14^C ages as individual Delta_R ‘likelihoods’, i.e. offsets from the atmospheric curve. The posterior distribution of each Delta_R indicates whether the estimated DRE is compatible with any dating constraints applied (e.g. the assumption that one sample is exactly contemporaneous with another). When using the diet‐reconstruction approach, ^14^C ages can also be calibrated with OxCal’s Mix_Curves function, which applies user‐defined mixtures to each ^14^C age, based on the FRUITS output, of the atmospheric calibration curve and calibration curve(s) for fish with a user‐defined Reservoir, i.e. offset from the atmospheric curve [e.g., 53,54]. In this case, chronological modelling provides posterior distributions for the curve mixture likelihoods, usually specified as normal distributions (mean ± 1σ). In ReSources (https://isomemoapp.com/app/resources), an online package using the same algorithms as FRUITS, the posterior distribution of the fish contribution to δ^13^C can be exported as a probability density function for the likelihood of the Mix_Curves function in OxCal.

Our Bayesian chronological model (Supplementary Information) incorporates several dating constraints, based on archaeological evidence or reasoning:

‐ Where an animal bone and one or more human bones from the same grave have been dated, we assume that they were exactly contemporaneous
‐ Where there are two or more AMS dates for the same individual, we assume that they refer to the same event (i.e. we regard the individual lifetime as trivial, relative to the uncertainty in calibrated date)
‐ We attribute all Lyalovo burials to a single period of burial activity, but have no prior information about the sequence of Lyalovo burials or their temporal distribution within this period
‐ We assume all Volosovo burials are later than all Lyalovo burials, but we do not a priori know the temporal distribution of Volosovo burials, which is one of the research questions
‐ We regard Volosovo burials and hoards at the four cemeteries as separate, potentially overlapping phases within the period of Volosovo burial activity
‐ We do not include the anomalous burials Sakhtysh IIa grave 11 and Sakhtysh II grave 12 in either period.

Our model does not incorporate:

‐ An assumption that Lyalovo burials postdate the early Neolithic Upper Volga culture occupation of Sakhtysh IIa, which may have lasted longer than previously suggested [10], due to the larger FRE implied by our results, although probably not beyond the end of the 6^th^ millennium
‐ An assumption that the Volosovo cultural layer at Sakhtysh IIa started before the first Volosovo burials, and ended before the last Volosovo burials; our results are compatible with this stratigraphic sequence, but the lack of precise dates for the cultural layer means that it provides no improvement in the burial chronology
‐ Any genetic kinship information, which would constrain potential differences in (birth) date between related individuals [e.g., 55], as this would require the integration of Kiel and Copenhagen ancient DNA results, which is not yet possible.

The model repeatedly samples all the likelihoods (e.g. probability distributions for calibrated dates), rejecting solutions that are incompatible with the ‘prior information’ (dating constraints), and retaining feasible solutions. After thousands of iterations, the cumulative distributions of feasible solutions provide posterior density estimates for the values of parameters such as the date of each burial, the first and last burial in each phase, and individual DREs. If nearly all the posterior distributions are consistent with the corresponding likelihoods, OxCal’s dynamic index of agreement A_model_ will be above a threshold value of 60. This does not mean that the model output is true, but it implies that there are no serious contradictions between archaeological reasoning and scientific data. We use the OxCal functions First, Last, Difference and Sum to summarise aspects of the model output (e.g. Figure 4).

## Acknowledgments

Analyses reported here were funded jointly by the Centre for Baltic and Scandinavian Archaeology, ZBSA research theme Man and Environment (Schleswig, Germany), the Graduate School Human Development and Landscapes (Christian‐Albrechts‐University Kiel, Germany), and sub‐cluster Dietary ROOTS of the ROOTS Cluster of Excellence (Christian‐Albrechts‐University Kiel, Germany). The material analysed for this study is curated with the support of the Center for Collective Use “Funds of Anthropological Materials of the Center of Physical Anthropology of the IEA RAS”, a grant provided by the Ministry of Science and Higher Education of the Russian Federation (grant ID: 075‐15‐2022‐ 328).

## Author contributions

Conceptualization: AK, HP, JM, BKK; Material preparation and data collection: JM, AK and BKK. Funding acquisition: JM, AK, HP, BKK. Resources: EK, EV, MD, SV. Supervision: HP. Original methodology: JM. Data analysis and investigation: JM, AK, BKK, NdS, HP; Writing ‐ original: JM; ‐ review and editing: JM, AK, EK, SV, BKK, NdS, HP. All authors read and approved the final manuscript.

## Data availability statement

Previously unpublished data are given in Table 1. Relevant published data which are necessary to reproduce our findings are reproduced, with appropriate citations, in Supplementary Tables 1 and 2.

