## Supplementary for "Dietary ^14^C reservoir effects and the chronology of prehistoric burials at Sakhtysh, central European Russia"

### Supplementary Information

#### Contents

S1. Quality assurance of new isotopic results 2

S2. Legacy isotopic data from prehistoric human remains at Sakhtysh 3

S3. Comparison of $\delta^{13}C$ and $\delta^{15}N$ results from different investigations 5

S4. Comparison of radiometric and AMS $^{14}C$ results on the same skeletons 7

S5. Comparison of AMS $^{14}C$ results on different elements of the same skeletons 8

S6. Comparison of human and osseous grave good $^{14}C$ ages from the same graves 9

S7. Intra-skeletal variation in AMS $^{14}C$ ages, $\delta^{13}C$ and $\delta^{15}N$ values 10

S8. DRE predictions from different MLR-of-difference formulae 11

S9. Faunal collagen $\delta^{13}C$ and $\delta^{15}N$ values 12

S10. OxCal model code 13

Sakhtysh OxCal model code 15

Fatyanovo burials OxCal model code 19

S11. Phasing Volosovo burials at Sakhtysh 23

S12. Carbon isotope systematics in freshwater ecosystems 24

S13. Comparison of DREs predicted by the MLR-of-differences and diet-reconstruction approaches 25

References 26

#### S1. Quality assurance of new isotopic results

All the dated extracts (Table 1) meet the usual standards for good collagen preservation, despite relatively low collagen yields (occasionally <1% of the starting weight) (Fig S1). %C and %N outliers are not associated with unusual isotope values or low collagen yields. Yields from samples extracted in Groningen (2.0±0.3%, n=10) are lower than those from Kiel (6.7±0.7%, n=22), presumably due in part to differences in extraction protocols. Among Groningen samples, collagen yield is not correlated with any other result (%C, %N, C/N, $\delta^{13}$C, $\delta^{15}$N, $^{14}$C age). Among Kiel samples, yields were highest in the Lyalovo cases, whose $^{14}$C ages and $\delta^{15}$N values are also higher, but among Volosovo individuals, yields are not correlated with other results. Thus the lower collagen yields do not appear to affect the new IRMS and AMS data.

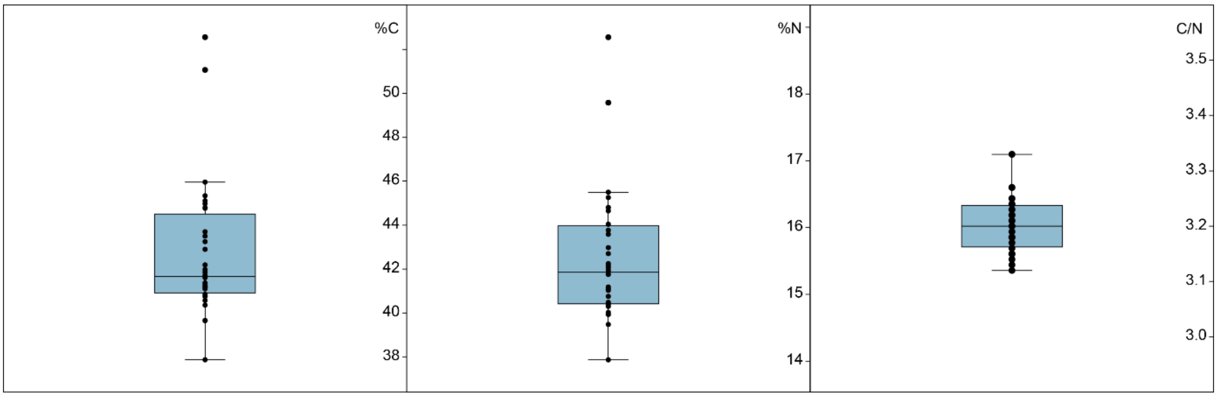


**Figure S1. Box-and-whisker/jitter plots of elemental-analyser results (Table 1) on newly dated collagen extracts (left, %C by weight; centre, %N by weight; right, atomic C/N value). Expected values for modern collagen are 41.91 ± 0.39 %C, 15.40 ± 0.20 %N, atomic C/N 3.00 to 3.28 [1]. The theoretical C/N value in unaltered human bone collagen is 3.243 [2].**

#### S2. Legacy isotopic data from prehistoric human remains at Sakhtysh

**Table S1: legacy data from Sakhtysh human bone/tooth samples (* $^{14}C$ age rejected by [3])**

| lab code | grave | phase | element | sex | age-at-death | $\delta^{13}C$ (‰) | $\delta^{15}N$ (‰) | $^{14}C$ age (BP) | source |
|---|---|---|---|---|---|---|---|---|---|
| GIN-6234 | IIa grave 10 | Volosovo | bone | F | 20–25 | | | 4540±160 | [3] |
| GIN-6237 | IIa grave 5 | Volosovo | bone | M | 40–45 | | | 4800±200 | [3] |
| GIN-6586 | IIa grave 42 | Lyalovo | bone | M | 20–25 | | | 6060±150 | [3] |
| GIN-6587 | IIa grave 43 | Lyalovo | bone | - | 5–6 | | | 8700±800* | [3] |
| GIN-7185 | IIa grave 12 | Lyalovo | bone | M | 30–40 | | | 6110±200 | [3] |
| GIN-7187 | IIa grave 14 | Volosovo | bone | M | c.40 | | | 5380±140* | [3] |
| GIN-7189 | IIa grave 13A | Volosovo | bone | F | 35–40 | | | 4200±240 | [3] |
| GIN-7190 | IIa grave 28 | Volosovo | bone | M | 35–40 | | | 4740±110 | [3] |
| GIN-7195 | IIa grave 29 | Lyalovo | bone | F | 40–45 | | | 5820±200 | [3] |
| GIN-7270 | IIa grave 36 | Volosovo | bone | M | 20–25 | | | 5090±90* | [3] |
| GIN-7271 | IIa grave 32B | Volosovo | bone | M | 40–45 | | | 3040±200* | [3] |
| GIN-7272 | IIa grave 36A | Volosovo | bone | M | 40–45 | | | 2030±260* | [3] |
| GIN-7273 | IIa grave 35 | Volosovo | bone | M | 35–40 | | | 4080±180 | [3] |
| GIN-7274 | IIa grave 32A | Volosovo | bone | M | - | | | 7730±70* | [3] |
| GIN-7275 | IIa grave 31 | Volosovo | bone | F | - | | | 5540±150* | [3] |
| GIN-7276 | IIa grave 34 | Volosovo | bone | M | 50–55 | | | 4540±150 | [3] |
| GIN-7277 | IIa grave 33 | Volosovo | bone | M | 50–55 | | | 3550±200* | [3] |
| GIN-7490 | IIa grave 64 | Volosovo | bone | F | 45–50 | | | 4550±350 | [3] |
| GIN-7492 | IIa grave 16 | Lyalovo | bone | F | 20–25 | | | 6130±120 | [3] |
| AAR-15050 | IIa grave 40 | Lyalovo | bone | M | 50–60 | -20.9 | 13.4 | 6406±24 | [4] |
| AAR-15052 | IIa grave 61 | Lyalovo | bone | F | 20–25 | -21.1 | 14.6 | 6356±23 | [4] |
| AAR-15051 | IIa grave 54 | Volosovo | bone | F | 45-50 | -21.4 | 12.4 | 4964±23 | [4] |
| AAR-15053 | IIa grave 66 | Volosovo | bone | F | 20–25 | -23.0 | 12.6 | 5033±24 | [4] |
| UBA-39973 | II grave 4 | Volosovo | tooth | M | | -21.6 | 13.8 | 5287±40 | [5] |
| UBA-39990 | II grave 12 | transitional | tooth | M | 30-35 | -21.8 | 12.4 | 4754±50 | [5] |
| UBA-39991 | IIa grave 13 sk.2 | Volosovo | tooth | F | 50–60 | -23 | 12.7 | 4919±36 | [5] |
| UBA-39992 | IIa grave 36 lower | Volosovo | tooth | M | mature | -23.2 | 12.7 | 5314±34 | [5] |
| UBA-39993 | IIa grave 58 | Volosovo | tooth | M | 45-50 | | | 5328±39 | [5] |
| UBA-39994 | IIa grave 46 | Volosovo | tooth | M | 25-30 | -22.5 | 12.5 | 4767±35 | [5] |
| UBA-39995 | IIa grave 33 | Volosovo | tooth | M | 50-55 | -23.4 | 13.4 | 5011±35 | [5] |
| UBA-39996 | VIII trench 1 | Volosovo | tooth | F | 30-35 | -23.1 | 13.3 | 5014±36 | [5] |
| UBA-39997 | IIa grave 40 | Lyalovo | tooth | M | 50-55 | -22.2 | 15.3 | 6393±39 | [5] |
| UBA-39998 | IIa grave 42 | Lyalovo | tooth | M | 30-35 | -22.6 | 14.3 | 6317±91 | [5] |
| UBA-39999 | IIa grave 11 | transitional | tooth | F | 20-25 | -22.4 | 13.9 | 4616±38 | [5] |
| UBA-40000 | IIa grave 9 | Volosovo | tooth | M | 50-55 | -23.3 | 13.9 | 4916±35 | [5] |
| UBA-40001 | IIa grave 39 | Volosovo | tooth | M | 30-35 | -24.6 | 14.6 | 5157±35 | [5] |
| UBA-40003 | II grave 19 | Lyalovo | tooth | F | 20-25 | -20.1 | 13.2 | 6265±38 | [5] |
| UBA-40004 | IIa grave 32 | Volosovo | tooth | M | 40-45 | -23.7 | 13.7 | 4981±37 | [5] |
| UBA-40005 | IIa grave 34 | Volosovo | tooth | M | 45-50 | -24.4 | 13.9 | 5143±34 | [5] |
| UBA-40006 | IIa grave 35 | Volosovo | tooth | M | 45-50 | -22.8 | 12.6 | 5118±59 | [5] |
| UBA-40007 | IIa grave 36 | Volosovo | tooth | M | 35-40 | -23.5 | 13.7 | 4827±34 | [5] |

**Table S2: IRMS results from Sakhtysh IIa human bones [6]**

| grave | archaeological phasing | age-at-death | δ$^{13}$C (‰) | δ$^{15}$N (‰) |
|---|---|---|---|---|
| grave 40 | Lyalovo | 40–49 | -20.9 | 13.4 |
| grave 42 | Lyalovo | 30–39 | -21.0 | 13.1 |
| grave 61 | Lyalovo | 10 | -20.6 | 14.0 |
| grave 16 | probably Lyalovo | 10 | -20.2 | 12.6 |
| grave 29 | probably Lyalovo | 5–9 | -17.7 | 12.1 |
| grave 5 (no.82) | Volosovo | 10–14 | -21.0 | 11.4 |
| grave 14 | Volosovo | 35–44 | -21.9 | 12.2 |
| grave 27 | Volosovo | 25–34 | -21.3 | 11.9 |
| grave 33 | Volosovo | 30–39 | -22.1 | 11.3 |
| grave 1 | probably Volosovo | 35–44 | -22.2 | 11.3 |
| grave 7 | probably Volosovo | 25–34 | -21.2 | 12.3 |
| grave 13 | probably Volosovo | 30–39 | -23.6 | 11.2 |
| grave 21 | probably Volosovo | 20–24 | -21.3 | 12.1 |
| grave 24 | probably Volosovo | 10–12 | -20.7 | 12.0 |
| grave 25 | probably Volosovo | 20–25 | -23.0 | 11.5 |
| grave 31 | probably Volosovo | 30–39 | -20.6 | 12.2 |
| grave 34 | probably Volosovo | 30–39 | -23.6 | 11.7 |
| grave 38 | probably Volosovo | 30–39 | -24.1 | 13.1 |
| grave 47 | probably Volosovo | 30–39 | -20.5 | 11.0 |
| grave 50 | probably Volosovo | 5–6 | -22.6 | 12.3 |
| grave 53 | unknown | 30–39 | -21.9 | 11.4 |
| grave 57 | unknown | adult | -22.9 | 12.7 |

#### S3. Comparison of $\delta^{13}$C and $\delta^{15}$N results from different investigations

There are now $\delta^{13}$C and $\delta^{15}$N data from 74 human samples, of which 52, representing at least 37 individuals, are also AMS-dated; none of these 52 gave an IRMS $\delta^{13}$C value above -20‰. The $\delta^{13}$C outlier, IIa grave 29 (-17.7‰) was replicated between laboratories in Moscow and St Petersburg [6], This burial has not been dated directly, and was unfurnished, apart from ochre, but the grave was aligned with other Lyalovo graves and was cut by Volosovo graves 18, 26 and 27 (the last of which we have dated, to the mid-4$^{th}$ millennium cal BC). Thus it is probably Lyalovo, but could be even older.

Adult bone $\delta^{13}$C values are generally higher, and/or $\delta^{15}$N values are lower, than those in petrous bones or teeth (Figure S2), leading to small differences in average $\delta^{13}$C and $\delta^{15}$N between our data and those of Engovatova et al. [6]. Comparing all Volosovo-transitional petrous bone results (mean $\delta^{13}$C -22.8±1.0‰, $\delta^{15}$N 12.7±0.8‰, n=27) to (probably) Volosovo bone data from [6] (mean $\delta^{13}$C -22.0±1.1‰, $\delta^{15}$N 11.9±0.6‰, n=17), both mean values are significantly different at the 5% significance level (2-tailed homoscedastic T-test). The higher petrous bone $\delta^{15}$N values cannot be attributed to a residual nursing effect [7], as this explanation would entail higher, not lower, $\delta^{13}$C values in petrous bones. The apparent dietary shift from childhood to adulthood would be more convincing if it was not visible in both Lyalovo and Volosovo samples, and if [6] had published quality-assurance data. We suspect that some adult bone extracts were affected by environmental contamination – not enough to obscure overall trends in $\delta^{13}$C and $\delta^{15}$N, but raising doubts about individual values. Adult bone $\delta^{13}$C, $\delta^{15}$N and $^{14}$C data reported by Piezonka [4] were supported by quality-assurance data and we regard them as accurate.

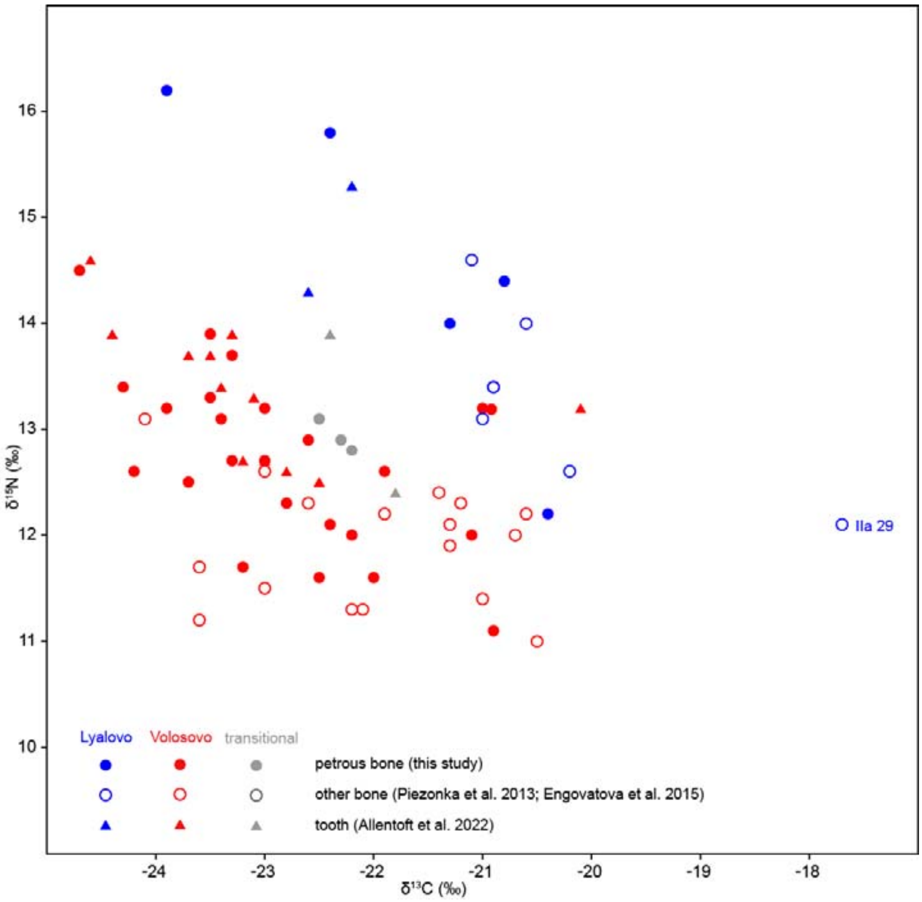


**Figure S2: IRMS $\delta^{13}$C and $\delta^{15}$N values from human remains, Sakhtysh prehistoric cemeteries (Table 1, Tables S1, S2). Symbols correspond to skeletal element sampled; colours represent chronological phase attribution (confirmed by AMS dating of petrous bones, teeth and some other bone samples, and stratigraphic position of IIa grave 29).**

Among the AMS-dated samples, there is a clear isotopic separation between periods, but in both periods the two isotopes appear to be correlated (Figure S3), as expected if variation is due mainly to differential consumption of two food groups with different $\delta^{13}C$ and $\delta^{15}N$ signatures. Our preferred explanation for the Lyalovo-Volosovo isotopic offset is the availability in the Lyalovo period of both high- and low-$\delta^{13}C$ fish (see Discussion and S12, Carbon isotope systematics in freshwater systems).

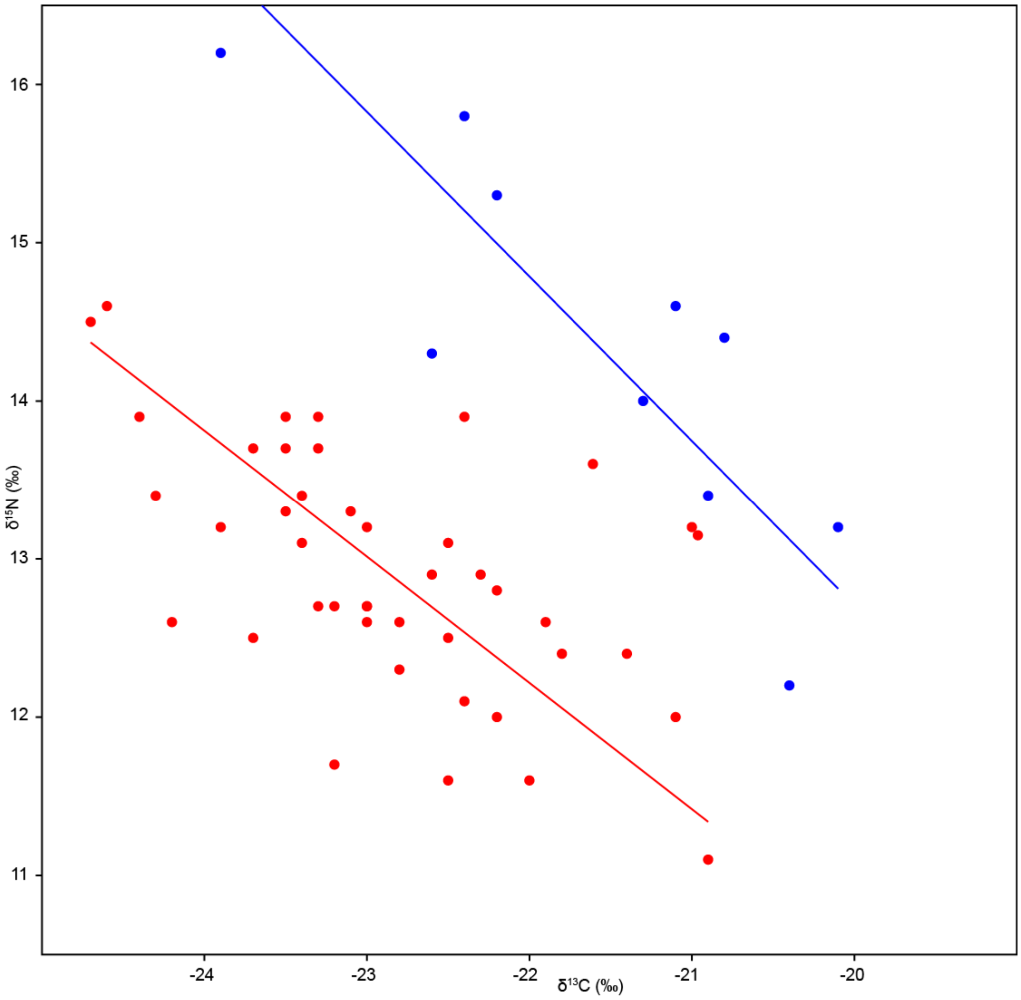


**Figure S3: Reduced Major Axis regressions of Lyalovo (blue) and Volosovo-transitional (red) $\delta^{13}C$ and $\delta^{15}N$ values in AMS-dated samples (mainly petrous bones and teeth; Tables 1 and S1)**

#### S4. Comparison of radiometric and AMS $^{14}C$ results on the same skeletons

We dated a petrous bone from nine of the individuals dated in the original radiometric dating programme [3]. Only two pairs of $^{14}C$ results (graves 5 and 42; Figure S4) are consistent at the 5% significance level [8]. In graves IIa 10 and 14, $^{14}C$ age offsets could be due to DRE differences between different skeletal elements, but in most cases the offsets are too large (700-3000 $^{14}C$ years). Up to three further cases were replicated by [5] (Table S1): IIa grave 34, and two individuals from IIa grave 36. In each case, the paired radiometric-AMS results are incompatible, although the 229±96 $^{14}C$ year difference between GIN-7270 and UBA-39992 could reflect diet changes between tooth formation and bone remodelling. Given this pattern, we have not attempted to include the radiometric results in the chronological model. At best, they would have provided maximum dates (without dietary stable isotopes on these extracts we cannot estimate DREs), but as some of the radiometric $^{14}C$ ages are clearly too recent, such maximum dates would not be reliable.

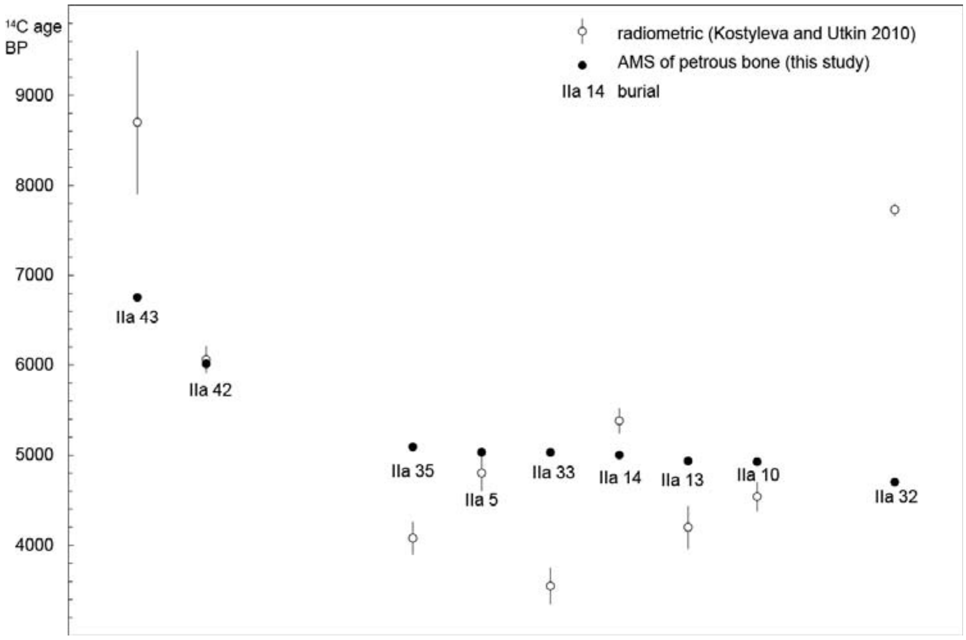


**Figure S4: comparison of AMS and radiometric $^{14}C$ ages (Table 1, Table S1) in cases where the same individual has been dated twice. Error bars (1σ) for AMS measurements are smaller than the symbols.**

#### S5. Comparison of AMS $^{14}C$ results on different elements of the same skeletons

In 8 of the 16 cases where we can compare AMS $^{14}C$ ages from different skeletal elements of the same individual, the differences are statistically significant (Figure S5). There is no consistent pattern of inter-laboratory offsets that might suggest analytical errors. Instead, we propose that the $^{14}C$ age differences correspond to differences in DRE between skeletal elements, due to differences in collagen formation time and lifetime dietary changes. Collagen turnover in petrous bone is negligible, which means that the isotopic results correspond to the diet in early childhood [9]. Allentoft et al. [5] do not report which teeth they sampled, but these samples probably represent short intervals of childhood or adolescence, which in many cases did not coincide with petrous bone collagen formation. Isotope values in bones sampled by Piezonka [4] represent long-term diets; the individuals concerned all died in adulthood, so these bone samples must have continued remodelling long after the petrous bones.

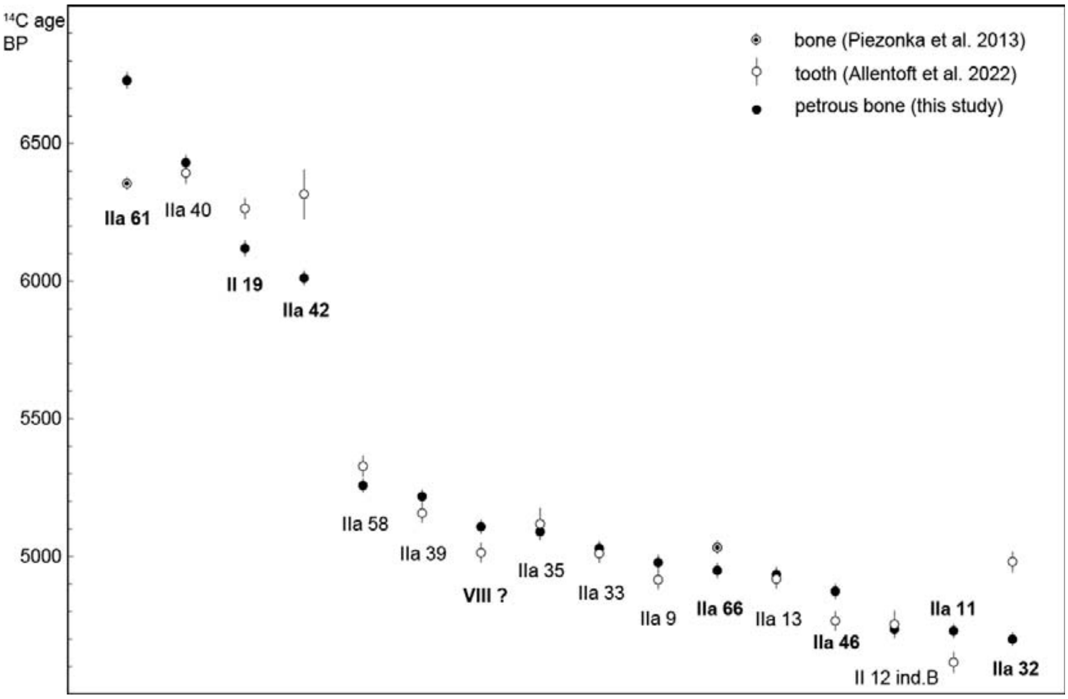


**Figure S5: comparison of AMS $^{14}C$ ages (1-σ error bars) from different skeletal elements of the same individual (Tables 1 and S1). Labels in bold type denote cases where $^{14}C$-age differences are statistically significant, following [8].**

#### S6. Comparison of human and osseous grave good $^{14}C$ ages from the same graves

Macāne et al. [10] dated osseous (bone or tooth) grave goods from five burials at Sakhtysh. In four cases, we sampled human petrous bones from the same burials (Table S3). In one case, the $^{14}C$ ages are not significantly different; in two cases, the human $^{14}C$ age is about 400 years older than the grave good; and in one case, the human $^{14}C$ ages are significantly younger. In this case, however, the bone beads are attributed to an aquatic species (fish, bird or otter), based on $\delta^{13}C$ and $\delta^{15}N$ values [10], so we assume that their $^{14}C$ ages incorporate larger DREs than those of the associated human remains. The alternative interpretation, that the beads predate the burial, is untenable considering the number of similar beads in this grave. The bear teeth $\delta^{13}C$ and $\delta^{15}N$ values provide no indication of fish consumption [10], so we assume that their $^{14}C$ ages are not subject to DREs. If bears consumed significant amounts of fish, the human-bear $^{14}C$ differences would under-estimate human DREs.

**Table S3. Comparison of human bone and osseous grave good $^{14}C$ ages.**

| grave | human petrous bone $^{14}C$ ages (this study) | osseous grave good $^{14}C$ ages [10] | difference (human - grave good) |
|---|---|---|---|
| IIa grave 19 | KIA-54657, 4931±29 | bear tooth UBA-34990, 4881±42 | 50±51 (1.0 σ) |
| IIa grave 7B | GrM-17396, 5145±25 | bear tooth UBA-34099, 4719±45 | 426±51 (8.4 σ) |
| IIa grave 24-25 | KIA-53532, 5177±25 | marmot tooth UBA-34098, 4769±38 | 408±45 (9.1 σ) |
| II grave 12 | ind. A KIA-53559, 4644±24<br>ind. B KIA-53560, 4736±25<br>human average 4689±18 | bone bead UBA-34097, 5170±40<br>bone bead UBA-40193, 5164±38<br>bead average 5167±28 | -478±33 (14.5 σ) |

The difference between two independent $^{14}C$ ages, $t_A \pm \sigma_A$ and $t_B \pm \sigma_B$, i.e. $t_{A-B} \pm \sigma_{A-B}$, is $(t_A - t_B) \pm \sqrt{(\sigma_A^2 + \sigma_B^2)}$. Two results are significantly different if the difference between them is more than twice the uncertainty in the difference, i.e. $|t_{A-B}|/\sigma_{A-B} > 2$.

#### S7. Intra-skeletal variation in AMS $^{14}C$ ages, $\delta^{13}C$ and $\delta^{15}N$ values

**Table S4. Comparison of human bone and tooth $\delta^{13}C$, $\delta^{15}N$ and $^{14}C$ ages. In most cases, the tooth or adult bone values (Table S1) were subtracted from the petrous bone value (Table 1). In two cases (marked §), the $\delta^{13}C$, $\delta^{15}N$ and DRE of IIa grave 19's petrous bone were subtracted from the corresponding values in the petrous bones shown (Table S3).**

| individual | $\delta^{13}C$ difference (‰) | $\delta^{15}N$ difference (‰) | $^{14}C$ age difference (years) | $^{14}C$ age difference (sigma) |
|---|---|---|---|---|
| II 12 ind.B | 0.52 | -0.47 | 18 | 0.32 |
| II 19 | 0.27 | 1.02 | 145 | 3.02 |
| IIa 7B§ | -0.13 | 2.13 | 376 | 5.22 |
| IIa 9 | -1.37 | 1.26 | -62 | -1.38 |
| IIa 11 | 0.10 | 0.78 | -114 | -2.53 |
| IIa 13 sk.2 | 0.27 | 0.02 | -16 | -0.35 |
| IIa 25§ | -2.32 | 0.57 | 358 | 5.26 |
| IIa 32 | -0.88 | 1.4 | 281 | 6.24 |
| IIa 33 | -0.38 | 0.65 | -19 | -0.44 |
| IIa 35 | 0.55 | -0.49 | 28 | 0.43 |
| IIa 39 | -0.28 | 1.28 | -61 | -1.39 |
| IIa 40 | -1.36 | 0.89 | -39 | -0.81 |
| IIa 42 | -1.26 | 0.3 | 305 | 3.21 |
| IIa 46 | 0.77 | -1.22 | -107 | -2.38 |
| IIa 61 | 1.38 | -1.16 | -374 | -9.84 |
| IIa 66 | -0.39 | -0.29 | 84 | 2.33 |
| VIII | -0.09 | 0.15 | -94 | -2.14 |

The difference between two independent $^{14}C$ ages, $t_A \pm \sigma_A$ and $t_B \pm \sigma_B$, i.e. $t_{A-B} \pm \sigma_{A-B}$, is $(t_A - t_B) \pm \sqrt{(\sigma_A^2 + \sigma_B^2)}$. Two results are significantly different if the difference between them is more than twice the uncertainty in the difference, i.e. $|t_{A-B}|/\sigma_{A-B} > 2$.

#### S8. DRE predictions from different MLR-of-difference formulae

Predicted DREs are calculated assuming human $\delta^{13}C$ and $\delta^{15}N$ values for a 100% terrestrial diet of -20‰ and 11.6‰, and the following formulae, given by multiple linear regression of $^{14}C$ differences against $\delta^{13}C$ and $\delta^{15}N$ differences in samples from the same individual:

- 17-case regression (no cases omitted) (adjusted $r^2$=0.33): DRE increases by 89±52 years for every 1‰ decrease in $\delta^{13}C$ ($r^2$=0.347; $p_{uncorr}$=0.108), and by 66±52 years for every 1‰ increase in $\delta^{15}N$ ($r^2$=0.291; $p_{uncorr}$=0.290); correlations between $\delta^{13}C$, $\delta^{15}N$, and $^{14}C$ differences could have arisen by chance;
- 14-case regression (omitting IIa graves 9, 39 and 40; see Results) (adjusted $r^2$=0.68): DRE increases by 130±41 years for every 1‰ decrease in $\delta^{13}C$ ($r^2$=0.623; $p_{uncorr}$=0.009) and 93±40 years for every 1‰ increase in $\delta^{15}N$ ($r^2$=0.505; $p_{uncorr}$=0.042); correlations between $^{14}C$ differences and both $\delta^{13}C$ and $\delta^{15}N$ differences cannot have arisen by chance;
- 9-case regression (omitting cases with statistically insignificant $^{14}C$ differences) (adjusted $r^2$=0.79): DRE increases by 140±42 years for every 1‰ decrease in $\delta^{13}C$ ($r^2$=0.656; $p_{uncorr}$=0.015) and 110±42 years for every 1‰ increase in $\delta^{15}N$ ($r^2$=0.547; $p_{uncorr}$=0.038).

In Figure S6, predicted DREs are plotted against the observed DREs (Table S3) in the three cases where this is possible. Not surprisingly, given the weak correlations, predicted DREs from the 17-case regression formula are the least accurate.

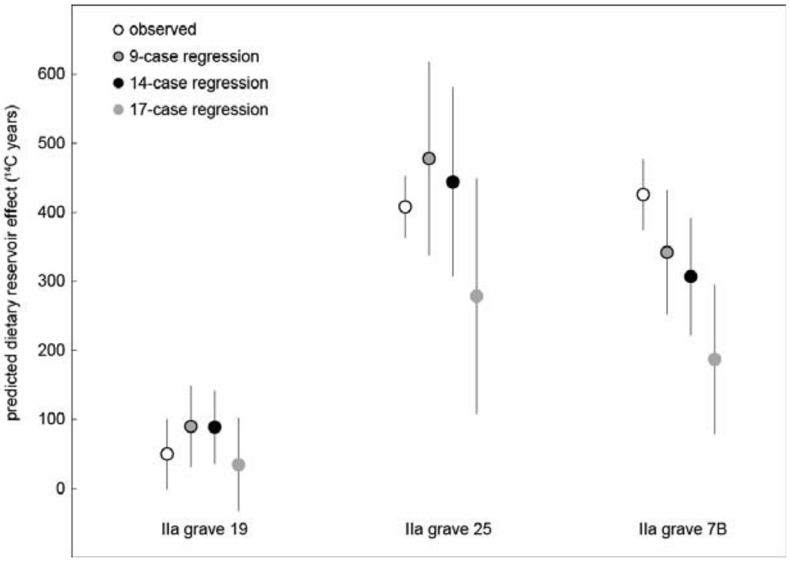


**Figure S6: comparison of $^{14}C$ age offsets between human petrous bones and herbivore teeth from the same burials and DREs in the same human bone samples predicted by multiple linear regression of $^{14}C$, $\delta^{13}C$ and $\delta^{15}N$ differences in 9 cases with significant $^{14}C$ age differences, in 14 cases, and in all 17 cases where results can be compared.**

Under a MLR approach, predicted DREs increase with isotopic distance from notional values for a fully terrestrial diet; DRE uncertainties are proportional to predicted DREs. The accuracy of predicted DREs depends on the regression slope, which is sensitive to which cases are omitted. Which cases are omitted also affects how noisy the regression is, and thus the relative uncertainty in predicted DREs. In the Sakhtysh case, predicted DREs would be smaller if all 17 cases were used in the MLR-of-differences, but their relative uncertainties would be larger. Consequently, the Volosovo phase would appear somewhat shorter, with no burials after c.3000 cal BC, and the temporal distribution of Volosovo burials would shift, with most potentially dating to the mid-4th millennium, and a tail of burials later in the 4th millennium. The 17-case regression is not satisfactory, however, as correlations between $^{14}C$ differences and $\delta^{13}C$ and $\delta^{15}N$ differences could have arisen by chance ($p_{uncorr}$>5% for both isotopes).

#### S9. Faunal collagen $\delta^{13}C$ and $\delta^{15}N$ values

Aside from 8 $^{14}C$ samples [4,10], only one faunal sample from Sakhtysh, an unidentified and undated bone from IIa grave 57, has published $\delta^{13}C$ and $\delta^{15}N$ values (-21.6‰ and 6.8‰) [6], which we do not use. Reference isotopic data from terrestrial faunal collagen including bones from Mesolithic-early Neolithic layers at Zamostje 2, 180 km west of Sakhtysh [11], Eneolithic-Bronze Age layers at Shagara, 150 km south of Sakhtysh [12], and Mesolithic layers at Minino I/II, 330 km north of Sakhtysh [13]. Together with the Sakhtysh data, these comprise 37 elks, 24 beavers, 2 wild boar and 1 marmot. Average $\delta^{13}C$ and $\delta^{15}N$ for all 64 animals (-22.0±0.5‰ and 5.1±1.2‰ respectively) are practically identical to average values for elk (-21.9±0.6‰ and 5.0±1.1‰). Elk $\delta^{13}C$ values at Shagara (mean -22.7±0.3‰, n=9) are somewhat lower than at the other sites, while the Zamostje elk $\delta^{15}N$ values (mean 5.8±0.8‰, n=16) are slightly higher than elsewhere, but these differences are relatively trivial, and the overall averages from all 64 animals are probably representative of terrestrial animals consumed at Sakhtysh.

Early Neolithic fish bones from Sakhtysh IIa did not yield collagen (Meadows unpublished data), and published data from comparable sites show contrasting patterns. At Shagara, 15 specimens of several fish taxa all gave lower $\delta^{13}C$ and higher $\delta^{15}N$ (mean -25.4±1.1‰ and 10.0±1.3‰) than terrestrial fauna. This pattern is probably typical of inland sites in the northeast European forest zone [14]. At Late Mesolithic Zamostje, however, both fish taxa sampled (cyprinids and pike) have $\delta^{15}N$ only slightly higher than elk (cyprinids 6.0±1.0‰, n=10, pike 6.9±0.6‰, n=10, and elk 5.8±0.8‰, n=16), while their mean $\delta^{13}C$ values are significantly different (cyprinids -25.5±1.3‰, pike -21.2±1.4‰). The much higher $\delta^{13}C$ values for young pike suggest that these fish were caught in a different water body to the cyprinids [11]. Zamostje may provide a better analogy than Shagara for the situation at Sakhtysh, for two reasons. One is the elevated $\delta^{13}C$, -17.7‰, for Sakhtysh IIa burial 29 [6], which cannot be explained by mixtures of terrestrial fauna with collagen $\delta^{13}C$ of c.-22‰ and fish with $\delta^{13}C$ of c.-25‰. Although not dated directly, this individual cannot date to a much later period when $C_4$ plants were available (see above). Some carbonised food crusts on early Neolithic pottery also have elevated $\delta^{13}C$ values [15], even higher than those of food crusts from Zamostje 2 [16]. Food crusts on two Volosovo sherds from Sakhtysh [10] have relatively low $\delta^{13}C$ values, however (-25 to -33‰).

During the Volosovo period, average fish collagen $\delta^{13}C$ must have been below c.-26‰ to account for human collagen $\delta^{13}C$ values approaching -25‰ (as at 4th-millennium Riņņukalns [14]). Modern fish from tributaries of the upper Volga in the Tver region have collagen $\delta^{13}C$ values of -31 to -28‰ [17], which, after correction for the Suess effect [e.g., 18], is compatible with the Riņņukalns data. In the Lyalovo period, average fish collagen $\delta^{13}C$ may have been 2-3‰ higher, given the human $\delta^{13}C$ and $\delta^{15}N$ values. The average $\delta^{15}N$ in fish collagen may also have varied between periods, as the maximum $\delta^{15}N$ in Lyalovo humans is 2‰ higher than the maximum Volosovo $\delta^{15}N$ value. Our diet reconstruction model assumes fish collagen $\delta^{13}C$ of -27±1‰ and $\delta^{15}N$ of 9.0±1‰ throughout the Volosovo period, and tests higher values of both isotopes in the Lyalovo period.

#### S10. OxCal model code

We use the program OxCal v4.4 (https://c14.arch.ox.ac.uk/oxcal/OxCal.html) for chronological modelling. Our Sakhtysh OxCal model (code below, to allow readers to reproduce the full output) calibrates human $^{14}C$ ages using the Delta_R function to apply the individual DRE estimated by the 14-case MLR-of-differences model to each result, and the Combine function to provide a single date for each burial with multiple $^{14}C$ measurements (including grave good dates where applicable, and of both humans in multiple burials). We use all the AMS dates from burial contexts, apart from those of two artefacts regarded as residual (see Introduction), which apparently pre-date the start of Volosovo burial activity. Our model omits radiometric $^{14}C$ ages on human bone and on bulk charcoal from cultural layers and sanctuary contexts, some already rejected by Kostyleva and Utkin [3] and Macāne et al. [10]. These $^{14}C$ measurements could be accurate, but bulk charcoal samples can include fragments with a range of different dates, giving misleading average dates. We also omit food-crust $^{14}C$ ages from two Volosovo potsherds, which may embody $^{14}C$ reservoir effects [10].

We use the KDE_Model function [19] to limit the dispersion of dates caused by measurement and calibration uncertainty, rather than imposing an arbitrary temporal distribution (e.g. assuming a uniform rate of burial), primarily because one of the research questions was whether Volosovo burial activity can be broken into two or more phases. As Lyalovo and Volosovo burials are clearly separable, even with the large dating uncertainties produced by DRE correction, we place them in separate KDE_Models. The OxCal functions First, Last, Difference and Sum are used to summarise aspects of the model output (e.g. Figure 4).

Our model has good overall and dynamic agreement ($A_{overall}$ = 90; $A_{model}$ = 121; a threshold value of 60 is considered adequate [20]), primarily because DRE-corrected calibrated dates of multiple samples from the same individual are generally compatible. In a few cases, the predicted DREs may be wrong, but there is no systematic pattern (Figure S7). As we omitted 3 of 17 potential cases from the MLR-of-differences regression, we may expect c.9 of the 53 predicted DREs to be misleading, but the OxCal model is not sufficiently informed by independent dating evidence to identify all of these cases.

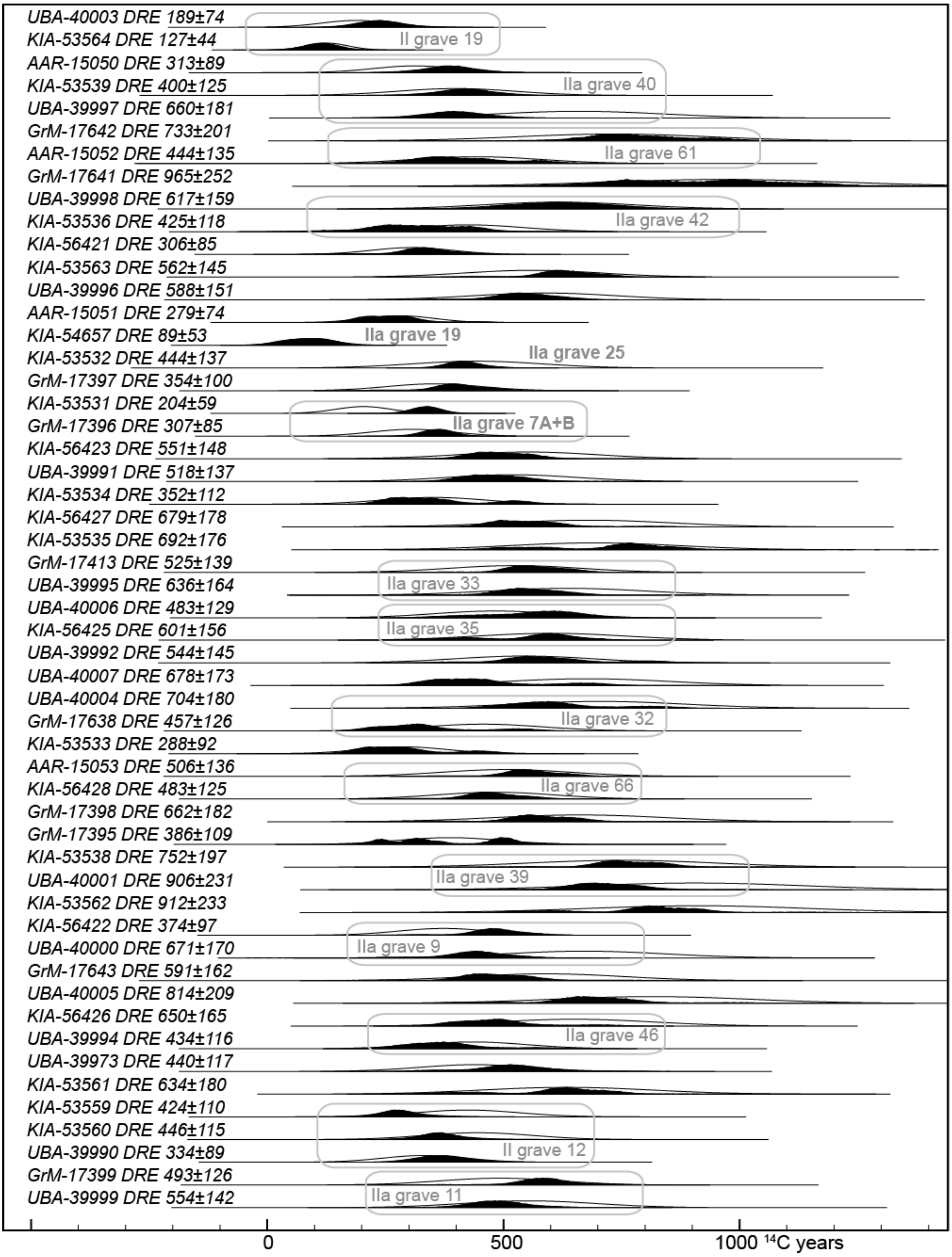


**Figure S7: comparison of predicted DREs in all AMS-dated human samples, 14-case MLR-of-differences approach (outline distributions) and posterior density estimates of these DREs from the Sakhtysh chronological model output (solid distributions). Burials with dated osseous grave goods (whose dates tightly constrain the posterior DRE estimates) are labelled in bold type. Other labels group multiple samples from the same grave.**

Meadows et al., *Dietary $^{14}$C reservoir effects and the chronology of prehistoric burials at Sakhtysh, central European Russia*. Supplementary Information.

#### Sakhtysh OxCal model code

Approximate geographical coordinates of Sakhtysh IIa graves are included to enable the OxCal Plot on Map function.

```
Plot()
{
 Curve("terrestrial","intcal20.14c");
 Sequence("Sakhtysh cemeteries")
 {
  //corrections of human 14C ages based on 14-case regression
  KDE_Model("Lyalovo burials (n=5)", )
  {
   Combine("II grave 19")
   {
    Delta_R("UBA-40003 DRE 189±74", 189, 74);
    R_Date("II grave 19 UBA-40003", 6265, 38);
    Delta_R("KIA-53564 DRE 127±44", 127, 44);
    R_Date("II grave 19 KIA-53564", 6120, 29);
   };
   Combine("IIa grave 40")
   {
    Delta_R("AAR-15050 DRE 313±89", 313, 89);
    R_Date("IIa grave 40 AAR-15050", 6406, 24);
    Delta_R("KIA-53539 DRE 400±125", 400, 125);
    R_Date("IIa grave 40 KIA-53539", 6432, 28);
    Delta_R("UBA-39997 DRE 660±181", 660, 181);
    R_Date("IIa grave 40 UBA-39997", 6393, 39);
   };
   Combine("IIa grave 61")
   {
    Delta_R("GrM-17642 DRE 733±201", 733, 201);
    R_Date("IIa grave 61 GrM-17642", 6730, 30);
    Delta_R("AAR-15052 DRE 444±135", 444, 135);
    R_Date("IIa grave 61 AAR-15052", 6356, 23);
   };
   Delta_R("GrM-17641 DRE 965±252", 965, 252);
   R_Date("IIa grave 43 GrM-17641", 6755, 35);
   Combine("II grave 42")
   {
    Delta_R("UBA-39998 DRE 617±159", 617, 159);
    R_Date("IIa grave 42 UBA-39998", 6317, 91);
    Delta_R("KIA-53536 DRE 425±118", 425, 118);
    R_Date("IIa grave 42 KIA-53536", 6012, 26);
   };
   First("first dated Lyalovo burial");
   Last("last dated Lyalovo burial");
  };
  Page( );
  Interval("Lyalovo-Volosovo hiatus");
  KDE_Model("all Volosovo burials (n=32)", )
  {
   Delta_R("KIA-56421 DRE 306±85", 306, 85);
   R_Date("I grave 8 KIA-56421", 4821, 28);
   Combine("VIII 1965 trench 1 grave ?")
   {
    Delta_R("KIA-53563 DRE 562±145", 562, 145);
    R_Date("VIII 1965 trench 1 grave ? KIA-53563", 5108, 26);
    Delta_R("UBA-39996 DRE 588±151", 588, 151);
    R_Date("VIII 1965 trench 1 grave ? UBA-39996", 5014, 36);
   };
   Phase("Sakhtysh IIa Volosovo phase")
   {
    Delta_R("AAR-15051 DRE 279±74", 279, 74);
```

```
R_Date("IIa grave 54 AAR-15051", 4964, 23)
{
 latitude=56.78585;
 longitude=40.44899;
};
Phase("Row G")
{
 Combine("IIa grave 19")
 {
  Curve("=terrestrial");
  R_Date("IIa grave 19 bear UBA-34990", 4881, 42);
  Delta_R("KIA-54657 DRE 89±53", 89, 53);
  R_Date("IIa grave 19 KIA-54657", 4931, 29);
  latitude=56.78576;
  longitude=40.44918;
 };
 Combine("IIa grave 25-24")
 {
  Delta_R("KIA-53532 DRE 444±137", 444, 137);
  R_Date("IIa grave 25 KIA-53532", 5177, 25);
  Curve("=terrestrial");
  R_Date("IIa grave 24 marmot UBA-34098", 4769, 38);
  latitude=56.78579;
  longitude=40.44914;
 };
 Delta_R("GrM-17397 DRE 354±100", 354, 100);
 R_Date("IIa grave 27 GrM-17397", 5165, 25)
 {
  latitude=56.78577;
  longitude=40.44916;
 };
};
Phase("Row V")
{
 Combine("IIa grave 7")
 {
  latitude=56.78563;
  longitude=40.44926;
  Delta_R("KIA-53531 DRE 204±59", 204, 59);
  R_Date("IIa grave 7A KIA-53531", 5147, 25);
  Delta_R("GrM-17396 DRE 307±85", 307, 85);
  R_Date("IIa grave 7B GrM-17396 ", 5145, 25);
  Curve("=terrestrial");
  R_Date("IIa grave 7B bear UBA-34099", 4719, 45);
 };
 Combine("IIa grave 13 sk.2")
 {
  latitude=56.7856;
  longitude=40.44929;
  Delta_R("KIA-56423 DRE 551±148", 551, 148);
  R_Date("IIa grave 13 KIA-56423", 4935, 28);
  Delta_R("UBA-39991 DRE 518±137", 518, 137);
  R_Date("II grave 13 sk.2 UBA-39991", 4919, 36);
 };
};
Phase("Row B")
{
 Delta_R("KIA-53534 DRE 352±112", 352, 112);
 R_Date("IIa grave 5 KIA-53534", 5032, 25)
 {
  latitude=56.7857;
  longitude=40.44928;
 };
};
```

```
Phase("Row D")
{
 Curve("=terrestrial");
 R_Date("IIa grave 63 bear UBA-34989", 4766, 51)
 {
  latitude=56.78609;
  longitude=40.44891;
 };
 Delta_R("KIA-56427 DRE 679±178", 679, 178);
 R_Date("IIa grave 56 KIA-56427", 4980, 28)
 {
  latitude=56.78601;
  longitude=40.44893;
 };
 Delta_R("KIA-53535 DRE 692±176", 692, 176);
 R_Date("IIa grave 58 KIA-53535", 5258, 26)
 {
  latitude=56.78598;
  longitude=40.44896;
 };
 Combine("IIa grave 33")
 {
  latitude=56.78589;
  longitude=40.44907;
  Delta_R("GrM-17413 DRE 525±139", 525, 139);
  R_Date("IIa grave 33 GrM-17413", 5030, 25);
  Delta_R("UBA-39995 DRE 636±164", 636, 164);
  R_Date("IIa grave 33 UBA-39995", 5011, 35);
 };
 Combine("IIa grave 35")
 {
  latitude=56.78588;
  longitude=40.44908;
  Delta_R("UBA-40006 DRE 483±129", 483, 129);
  R_Date("IIa grave 35 UBA-40006", 5118, 59);
  Delta_R("KIA-56425 DRE 601±156", 601, 156);
  R_Date("IIa grave 35 KIA-56425", 5090, 28);
 };
 Delta_R("UBA-39992 DRE 544±145", 544, 145);
 R_Date("IIa grave 36 lower UBA-39992", 5314, 34)
 {
  latitude=56.78587;
  longitude=40.44909;
 };
 Delta_R("UBA-40007 DRE 678±173", 678, 173);
 R_Date("IIa grave 36 UBA-40007", 4827, 34)
 {
  latitude=56.78587;
  longitude=40.44909;
 };
 Combine("IIa grave 32")
 {
  latitude=56.7859;
  longitude=40.44906;
  Delta_R("UBA-40004 DRE 704±180", 704, 180);
  R_Date("IIa grave 32 UBA-40004", 4981, 37);
  Delta_R("GrM-17638 DRE 457±126", 457, 126);
  R_Date("IIa grave 32 GrM-17638", 4700, 25);
 };
};
Phase("Row Zh")
{
 Delta_R("KIA-53533 DRE 288±92", 288, 92);
 R_Date("IIa grave 67 KIA-53533", 4967, 29)
```

```
  {
   latitude=56.78615;
   longitude=40.44887;
  };
  Combine("IIa grave 66")
  {
   latitude=56.78613;
   longitude=40.44891;
   Delta_R("AAR-15053 DRE 506±136", 506, 136);
   R_Date("IIa grave 66  AAR-15053", 5033, 24);
   Delta_R("KIA-56428 DRE 483±125", 483, 125);
   R_Date("IIa grave 66 KIA-56428", 4949, 27);
  };
  Delta_R("GrM-17398 DRE 662±182", 662, 182);
  R_Date("IIa grave 62 GrM-17398", 5035, 25)
  {
   latitude=56.78608;
   longitude=40.44897;
  };
 };
 Phase("Row A")
 {
  Delta_R("GrM-17395 DRE 386±109", 386, 109);
  R_Date("IIa grave 14 GrM-17395", 5000±25)
  {
   latitude=56.78556;
   longitude=40.4493;
  };
  Combine("IIa grave 39")
  {
   latitude=56.78577;
   longitude=40.44906;
   Delta_R("KIA-53538 DRE 752±197", 752, 197);
   R_Date("IIa grave 39 KIA-53538", 5218, 26);
   Delta_R("UBA-40001 DRE 906±231", 906, 231);
   R_Date("IIa grave 39 UBA-40001", 5157, 35);
  };
  Delta_R("KIA-53562 DRE 912±233", 912, 233);
  R_Date("IIa grave 15 KIA-53562", 5293, 26)
  {
   latitude=56.78569;
   longitude=40.44919;
  };
  Combine("IIa grave  9")
  {
   latitude=56.78562;
   longitude=40.44923;
   Delta_R("KIA-56422 DRE 374±97", 374, 97);
   R_Date("IIa grave 9 KIA-56422", 4978, 28);
   Delta_R("UBA-40000 DRE 671±170", 671, 170);
   R_Date("IIa grave 9 UBA-40000", 4916, 35);
  };
  Delta_R("GrM-17643 DRE 591±162", 591, 162);
  R_Date("IIa grave 10 GrM-17643", 4925, 30)
  {
   latitude=56.78558;
   longitude=40.44929;
  };
 };
 Phase("Row E")
 {
  Delta_R("UBA-40005 DRE 814±209", 814, 209);
  R_Date("IIa grave 34 UBA-40005", 5143, 34)
  {
```

```
      latitude=56.78592;
      longitude=40.44906;
     };
     Combine("IIa grave 46")
     {
      latitude=56.78595;
      longitude=40.44901;
      Delta_R("KIA-56426 DRE 650±165", 650, 165);
      R_Date("IIa grave 46 KIA-56426", 4874, 28);
      Delta_R("UBA-39994 DRE 434±116", 434, 116);
      R_Date("IIa grave 46 UBA-39994", 4767, 35);
     };
    };
   };
   Phase("Sakhtysh II Volosovo phase")
   {
    Delta_R("UBA-39973 DRE 440±117", 440, 117);
    R_Date("II grave 4 UBA-39973", 5287, 40);
    Delta_R("KIA-53561 DRE 634±180", 634, 180);
    R_Date("II grave 15 ind.7 KIA-53561", 5114, 26);
    Curve("=terrestrial");
    R_Date("II grave 18 bear UBA-40192", 4737, 31);
    R_Date("II hoard 9 bear UBA-34992", 4730, 41);
    R_Date("II hoard 11 badger UBA-34096", 4445, 37);
   };
   Last("last dated Volosovo burial");
   First("first dated Volosovo burial");
  };
 };
 Phase("transitional")
 {
  Combine("II grave 12")
  {
   Delta_R("KIA-53559 DRE 424±110", 424, 110);
   R_Date("II grave 12 ind.A KIA-53559", 4644, 24);
   Delta_R("KIA-53560 DRE 446±115", 446, 115);
   R_Date("II grave 12 ind.B KIA-53560", 4736, 25);
   Delta_R("UBA-39990 DRE 334±89", 334, 89);
   R_Date("II grave 12 UBA-39990", 4754, 50);
   Delta_R("bead FRE", 800, 100);
   R_Date("bead burial 12 UBA-40193", 5164, 38);
   R_Date("bead burial 12 UBA-34097", 5170, 40);
  };
  Combine("IIa grave 11")
  {
   latitude=56.78564;
   longitude=40.44929;
   Delta_R("GrM-17399 DRE 493±126", 493, 126);
   R_Date("IIa grave 11 GrM-17399", 4730, 25);
   Delta_R("UBA-39999 DRE 554±142", 554, 142);
   R_Date("IIa grave 11 UBA-39999", 4616, 38);
  };
 };
 Difference("duration of Lyalovo burials", "last dated Lyalovo burial", "first dated Lyalovo burial");
 Difference("duration of Volosovo burials", "last dated Volosovo burial", "first dated Volosovo burial");
 Difference("hiatus Lyalovo-Volosovo burials", "first dated Volosovo burial", "last dated Lyalovo burial");
};
```

#### Fatyanovo burials OxCal model code

Data from [21]. The model makes no allowance for possible dietary reservoir effects (DREs), which means that the estimated date of the first Fatyanovo burial ( First("Fatyanovo first");) is effectively a maximum age; any DREs, particularly among the earlier Fatyanovo burials, would mean that the first Fatyanovo burial was later than indicated by this model. Geographical coordinates are included to enable the OxCal Plot on Map function.

```
Plot()
 {
  KDE_Model("Saag 2021 dates",)
  {
   R_Date("VOR004 Voronkovo UBA-41640",4153,33)
   {
    latitude=57.5345164692551;
    longitude=39.5525236702395;
   };
   R_Date("NIK005 Nikultsino UBA-41629",4148,49)
   {
    latitude=57.529263726258;
    longitude=39.669940550546;
   };
   R_Date("NIK004 Nikultsino UBA-41628",4141,33)
   {
    latitude=57.529263726258;
    longitude=39.669940550546;
   };
   R_Date("NIK002 Nikultsino UBA-41626",4100,34)
   {
    latitude=57.529263726258;
    longitude=39.669940550546;
   };
   R_Date("IVA001 Ivanovogorsky UBA-41621",4092,35)
   {
    latitude=55.8225671366697;
    longitude=36.0651811705122;
   };
   R_Date("HAN002 Khanevo UBA-41618",4083,33)
   {
    latitude=55.63639;
    longitude=35.89972;
   };
   R_Date("NIK007 Nikultsino UBA-41630",4047,54)
   {
    latitude=57.529263726258;
    longitude=39.669940550546;
   };
   R_Date("NAU001 Naumovskoye UBA-41624",4047,35)
   {
    latitude=57.5870022863107;
    longitude=39.6308015904438;
   };
   R_Date("NIK008B Nikultsino UBA-41632 ",4039,34)
   {
    latitude=57.529263726258;
    longitude=39.669940550546;
   };
   R_Date("NIK008A Nikultsino UBA-41631 ",4039,33)
   {
    latitude=57.529263726258;
    longitude=39.669940550546;
   };
   R_Date("HAL001 Khaldeevo UBA-41617",4037,32)
```

```
{
 latitude=57.3205139320694;
 longitude=39.6216283966699;
};
R_Date("NAU002 Naumovskoye UBA-41625",4036,40)
{
 latitude=57.5870022863107;
 longitude=39.6308015904438;
};
R_Date("HAN004 Khanevo UBA-41619",4036,37)
{
 latitude=55.63639;
 longitude=35.89972;
};
R_Date("TIM008 Timofeyevka UBA-41637",4036,32)
{
 latitude=57.1344739083732;
 longitude=39.9757136763441;
};
R_Date("MIL001 Miloslavka UBA-41622",4033,29)
{
 latitude=56.9271425697216;
 longitude=39.6454787004822;
};
R_Date("TIM006 Timofeyevka UBA-41635",4031,36)
{
 latitude=57.1344739083732;
 longitude=39.9757136763441;
};
R_Date("BOL001 Bolshnevo 3 UBA-41613",4005,39)
{
 latitude=57.7400894913881;
 longitude=36.6330018651193;
};
R_Date("VOR005 Voronkovo UBA-41641",4002,54)
{
 latitude=57.5345164692551;
 longitude=39.5525236702395;
};
R_Date("VOR003 Voronkovo UBA-41639",3987,29)
{
 latitude=57.5345164692551;
 longitude=39.5525236702395;
};
R_Date("NIK003 Nikultsino UBA-41627",3972,54)
{
 latitude=57.529263726258;
 longitude=39.669940550546;
};
R_Date("GOL001 Goluzinovo UBA-41616",3968,32)
{
 latitude=57.3274474290247;
 longitude=39.5525236702395;
};
R_Date("BOL003 Bolshnevo 3 UBA-41615",3956,34)
{
 latitude=57.7400894913881;
 longitude=36.6330018651193;
};
R_Date("VOD001 Volosovo-Danilovsky UBA-41638",3943,41)
{
 latitude=57.8873303499581;
 longitude=40.2765944321294;
};
```

```
  R_Date("RDT002 Nikolo-Perevoz UBA-41634 ",3931,35)
  {
   latitude=56.533289621258;
   longitude=38.0915396751763;
  };
  R_Date("BOL002 Bolshnevo 3 UBA-41614",3876,36)
  {
   latitude=57.7400894913881;
   longitude=36.6330018651193;
  };
  R_Date("MIL002 Miloslavka UBA-41623",3763,30)
  {
   latitude=56.9271425697216;
   longitude=39.6454787004822;
  };
  First("Fatyanovo first");
 };
};
```

#### S11. Phasing Volosovo burials at Sakhtysh

Although typo-chronological schemes for Volosovo pottery and stone tools have been proposed, they cannot be used to phase the Sakhtysh burials, which did not contain diagnostic artefact types. Our OxCal model output suggests that there were two peaks of Volosovo burial activity, in the mid- and late 4th millennium cal BC (Figure 4b). Median modelled dates of individual burials (Figure 4b) are potentially misleading, however, due to the large uncertainties in DRE-corrected dates, particularly when only one $^{14}C$ date is available for a grave. In cases with mainly aquatic diets, such DRE-corrected dates can span >400 years at 95% probability. A more reliable approach is to use the posterior probability density functions in OxCal output, which provide a probability that each grave dates to every 5-year interval. We calculated the cumulative probability that each grave dates to between 3700 and 3400 cal BC, and the cumulative probability that the same grave dates to 3300-2900 cal BC (Figure S8). This separates the graves into two phases we have called earlier and later Volosovo. Three Volosovo graves could belong to either phase, or fall between these phases, as the cumulative probability of dating to one phase is <3× the cumulative probability of dating to the other phase (i.e. there is a >1 in 4 chance that if assigned to either earlier or later Volosovo phases, a grave would be assigned to the wrong phase). The anomalous multiple burial II grave 12 dates to the end of the later Volosovo phase, while the crouched burial IIa grave 11 is clearly later.

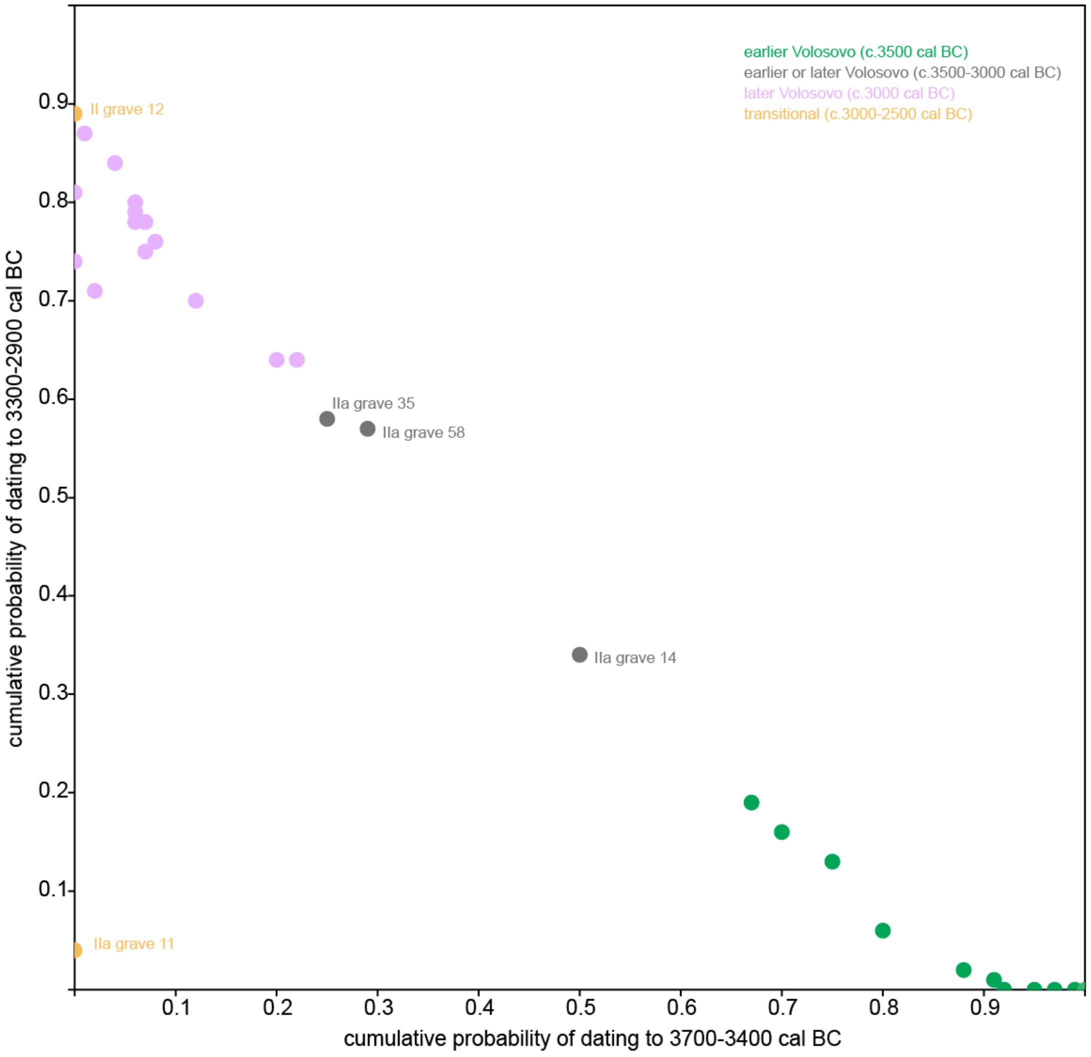


**Figure S8: separation of Volosovo burials into phases, based on the relative cumulative probability, given by the Sakhtysh OxCal chronological model output, that each grave dates to within the intervals 3700-3400 cal BC (horizontal axis) and 3300-2900 cal BC (vertical axis). For clarity, only graves which do not clearly belong to either earlier or later Volosovo phases are labelled (see text).**

#### S12. Carbon isotope systematics in freshwater ecosystems

One plausible explanation for the systematic difference in Lyalovo and Volosovo stable isotope values is that high-$\delta^{13}$C fish was more readily available in the earlier period, due to a change in local hydrology (e.g. if shallow lakes were infilled with peat) between the Lyalovo and Volosovo periods. There are indirect indications that high-$\delta^{13}$C fish was available in the early Neolithic (6th millennium cal BC), from the elevated $\delta^{13}$C values of some carbonised food crusts with aquatic biomarkers [15].

Freshwater fish in lakes and rivers of the northeast European forest zone typically have more negative collagen $\delta^{13}$C values than terrestrial mammals [e.g., 14], but in southern Siberia there are examples of higher $\delta^{13}$C values in fish from shallow lakes, in a region where river fish have low $\delta^{13}$C values like those in northeastern Europe [22]. In cold running water, primary productivity is limited by temperature and light, not the availability of dissolved inorganic carbon (DIC, specifically dissolved carbon dioxide, $CO_2^-$) [23]. Photosynthate $\delta^{13}$C values should therefore depend primarily on the mass balance between $CO_2^-$ derived from the atmosphere (-7‰), the decomposition of ($C_3$) vegetation (<-25‰), and isotopic exchange with dissolved carbonates ($HCO_3^-$, c.-14‰) [24,25]. Allowing for natural fractionation between $CO_2^-$ and photosynthates, $\delta^{13}$C values at the base of the aquatic food chain should thus be more negative than in terrestrial food chains. In clear still water, rapid plant growth in spring can deplete $CO_2^-$ levels, forcing primary producers to reduce fractionation, thereby increasing $\delta^{13}$C throughout the aquatic food web [23]. In shallow water, the proportion of $CO_2^-$ coming from atmospheric $CO_2$ is expected to be greater, further increasing $\delta^{13}$C values. This is partly because the water surface-to-volume ratio is greater, but the mechanism which probably explains elevated $\delta^{13}$C values here and in Siberia is seasonal productivity: by fixing any available $CO_2^-$, algal blooms reduce the $CO_2^-$ pressure, allowing more atmospheric $CO_2$ molecules to 'invade' the water.

Initially, we proposed two isotopically distinct aquatic food resources. Their isotopic signatures were inevitably arbitrary, but realistic in view of the patterns observed in human and food-crust $\delta^{13}$C values at Sakhtysh. Low-$\delta^{13}$C fish was assumed to have had an average collagen $\delta^{13}$C value of -27.0±0.5‰ and $\delta^{15}$N of 9.0±1.0‰ (higher than the Zamostje 2 cyprinids discussed in S9, which are lower trophic-level fish, as otherwise it would be impossible to explain human $\delta^{15}$N values as high as 16‰). High-$\delta^{13}$C fish was assumed to have had an average collagen $\delta^{13}$C value of -18.0±0.5‰ and $\delta^{15}$N of 7.0±1.0‰, like the young pike from Zamostje 2. Due to time-averaging, only lower trophic-level young fish would show a strong seasonal $\delta^{13}$C signal in collagen. Fish flesh $\delta^{13}$C values are likely to have changed more rapidly than collagen $\delta^{13}$C [25], and the collagen $\delta^{13}$C value of -18.0±0.5‰ may therefore under-estimate the food $\delta^{13}$C value in high $\delta^{13}$C fish.

Siberian high-$\delta^{13}$C fish have low or negligible FREs, in contrast to low $\delta^{13}$C fish in the same geological setting [22]. This is consistent with the idea that elevated $\delta^{13}$C values are not simply due to reduced fractionation during photosynthesis, but also reflect a much greater contribution of atmospheric $CO_2$ to the $CO_2^-$ pool, which can occur in summer due to reduced inflows of fresh water, warmer temperatures and longer hours of sunlight permitting rapid plant growth that metabolises all the available $CO_2^-$. Applying the same reasoning at Sakhtysh, we would expect a much lower FRE in high-$\delta^{13}$C fish than in low-$\delta^{13}$C fish. Thus the average FRE in fish consumed was probably lower during the Lyalovo period than during the Volosovo period, not higher as suggested if we use the same FRUITS parameters in both periods (see Discussion and S13).

#### S13. Comparison of DREs predicted by the MLR-of-differences and diet-reconstruction approaches

For sensitivity analyses, we compare the mean DREs predicted by the 14-case MLR-of-differences model for all 53 human samples with AMS $^{14}C$ ages and $\delta^{13}C$ and $\delta^{15}N$ to the DRE predicted by the FRUITS model output (median fish contribution to collagen $\delta^{13}C$ × average FRE in fish) over a range of potential FRE values. At an average FRE of 960 years, DREs for Volosovo samples predicted by the two approaches are essentially identical (Figure S7).

Using the same FRUITS parameters, Lyalovo cases fall below the Volosovo regression line, suggesting either that average FRE was higher in the Lyalovo period, or that the FRUITS model under-estimates Lyalovo fish consumption. Increasing Lyalovo fish $\delta^{13}C$ and $\delta^{15}N$ by 1‰ each, would reconcile Lyalovo diet-reconstruction and MLR-of-differences DRE estimates using the same 960-year FRE, an interpretation more consistent with differences in fishing technology between the Lyalovo and Volosovo periods than with environmental change. We prefer the second option, however (see S12). As a rough approximation, if there was a 10‰ difference between high-$\delta^{13}C$ fish and low-$\delta^{13}C$ fish protein, a 2‰ higher average fish $\delta^{13}C$ in the Lyalovo period implies that 20% of Lyalovo fish protein was from high-$\delta^{13}C$ fish. Increasing average fish $\delta^{13}C$ by 2‰ in the FRUITS model would reconcile Lyalovo MLR-of-differences and FRUITS-derived predicted DREs using an average FRE of c.800 years, compared to the 960 years providing the best fit between predicted DREs in the Volosovo period. This is consistent with an average FRE in high-$\delta^{13}C$ fish of 150 years, which is realistic in view of our understanding of carbon isotope systematics in freshwater ecosystems (S12).

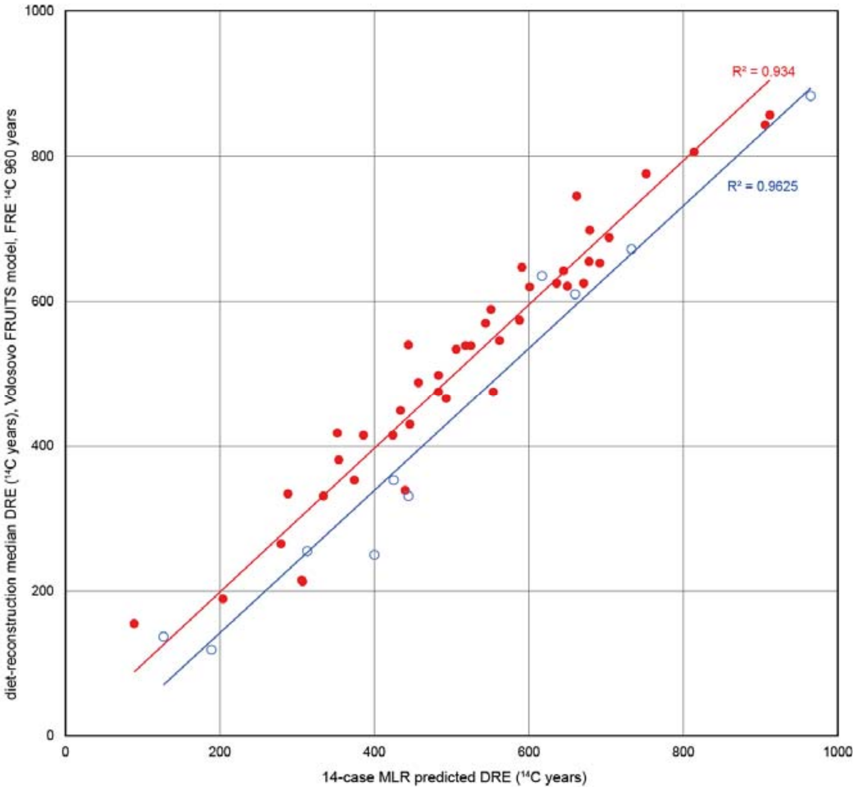


**Figure S7: comparison of predicted DREs in all AMS-dated human samples, 14-case MLR-of-differences approach (horizontal axis) vs diet-reconstruction approach (vertical axis), assuming an average 960-year FRE in local fish. For Volosovo-transitional samples (red dots), the two methods are perfectly compatible: a regression line with intercept set to zero has a slope of 0.995 and $r^2$ 0.9357, p<<0.001. For Lyalovo cases (blue circles), a regression line with a slope of 1.001 and $r^2$ 0.9625, p<<0.001, has an intercept of -63, i.e. the diet-reconstruction approach systematically under-estimates DREs compared to the MLR-of-differences approach if the FRE is set to 960 years.**
